# Deep ancestral introgression shapes evolutionary history of dragonflies and damselflies

**DOI:** 10.1101/2020.06.25.172619

**Authors:** Anton Suvorov, Celine Scornavacca, M. Stanley Fujimoto, Paul Bodily, Mark Clement, Keith A. Crandall, Michael F. Whiting, Daniel R. Schrider, Seth M. Bybee

## Abstract

Introgression is arguably one of the most important biological processes in the evolution of groups of related species, affecting at least 10% of the extant species in the animal kingdom. Introgression reduces genetic divergence between species, and in some cases can be highly beneficial, facilitating rapid adaptation to ever-changing environmental pressures. Introgression also significantly impacts inference of phylogenetic species relationships where a strictly binary tree model cannot adequately explain reticulate net-like species relationships. Here we use phylogenomic approaches to understand patterns of introgression along the evolutionary history of a unique, non-model insect system: dragonflies and damselflies (Odonata). We demonstrate that introgression is a pervasive evolutionary force across various taxonomic levels within Odonata. In particular, we show that the morphologically “intermediate” species of Anisozygoptera (one of the three primary suborders within Odonata besides Zygoptera and Anisoptera), which retain phenotypic characteristics of the other two suborders, experienced high levels of introgression likely coming from zygopteran genomes. Additionally, we found evidence for multiple cases of deep inter-superfamilial ancestral introgression.

## INTRODUCTION

In recent years there has been a large body of evidence amassed demonstrating that multiple parts of the Tree of Life did not evolve according to a strictly bifurcating phylogeny [1, 2]. Instead, many organisms experience reticulate network-like evolution that is caused by an exchange of inter-specific genetic information via various biological processes. In particular, lateral gene transfer, incomplete lineage sorting (ILS) and introgression can result in gene trees that are discordant with the species tree [3, 4]. Lateral transfer and introgression both involve gene flow following speciation, thereby producing “reticulate” phylogenies. Incomplete lineage sorting (ILS), on the other hand, occurs when three or more lineages coalesce in their ancestral population. This often arises in an order that conflicts with that of the true species tree, and thus involves no post-speciation gene flow and therefore does not contribute to reticulate evolution. Phylogenetic species-gene tree incongruence observed in empirical data can provide insight into underlying biological factors that shape the evolutionary trajectories of a set of taxa. The major source of reticulate evolution for eukaryotes is introgression where it affects approximately 25% of flowering plant and 10% of animal species [1, 5]. Introgressed alleles can be fitness-neutral, deleterious [6] or adaptive [7, 8]. For example, adaptive introgression has been shown to provide an evolutionary rescue from polluted habitats in gulf killifish (*Fundulus grandis*) [7], yielded mimicry adaptations among *Heliconius* butterflies [9] and archaic introgression has facilitated adaptive evolution of altitude tolerance [10], immunity and metabolism in modern humans [11]. Additionally, hybridization and introgression are important and often overlooked mechanisms of invasive species establishment and spread [12].

Odonata, the insect order that contains dragonflies and damselflies, lacks a strongly supported backbone tree to clearly resolve higher-level phylogenetic relationships [13, 14]. Current evidence places odonates together with Ephemeroptera (mayflies) as the living representatives of the most ancient insect lineages to have evolved wings and active flight [15]. Odonates possess unique anatomical and morphological features such as a specialized body form, specialized wing venation, a distinctive form of muscle attachment to the wing base [16] allowing for direct flight and accessory (secondary) male genitalia that support certain unique behaviors (e.g., sperm competition). They are among the most adept flyers of all animals and are exclusively carnivorous insects relying primarily on vision [17, 18] to capture prey. During their immature stage they are fully aquatic and spend much of their adult life in flight.

Biogeographically, odonates exhibit species ranges varying from worldwide dispersal [19] to island-endemic. Odonates also play crucial ecological roles in local freshwater communities, being a top invertebrate predator as both adults and immatures [20]. Due to this combination of characteristics, odonates are quickly becoming model organisms to study specific questions in ecology, physiology and evolution [21, 22]. However, the extent of introgression at the genomic scale within Odonata remains largely unknown.

Two early attempts to tackle introgression/hybridization patterns within Odonata were undertaken in [23, 24]. The studies showed that two closely related species of damselflies, *Ischnura graellsii* and *I. elegans*, can hybridize under laboratory conditions and that genital morphology of male hybrids shares features with putative hybrids from *I. graellsii*–*I. elegans* natural allopatric populations [23]. The existence of abundant hybridization and introgression in natural populations of *I. graellsii* and *I. elegans* has received further support from an analysis of microsatellite data [25]. Putative hybridization events have also been identified in a pair of calopterygoid damselfly species, *Mnais costalis* and *M. pruinosa* based on the analyses of two molecular loci (mtDNA and nucDNA) [26], and between *Calopteryx virgo* and *C. splendens* using 16S ribosomal DNA and 40 random amplified polymorphic DNA (RAPD) markers [27]. A more recent study identified an inter-specific hybridization between two cordulegasterid dragonfly species, *Cordulegaster boltonii* and *C. trinacriae* using two molecular markers (mtDNA and nucDNA) and geometric morphometrics [28]. Furthermore, in various biological systems, the empirical evidence shows that hybridization can potentially lead to intermediate phenotypes [29] observed at molecular level (e.g. semidominant expression in inter-specific hybrids [30]) as well as organismal morphology (e.g. [31-33]). The Anisozygoptera suborder, which contains only three extant species, retains traits shared with both dragonflies and damselflies (hence its taxonomic name), ranging from morphology and anatomical structures [34] to behavior and flight biomechanics [35]. These characteristics could suggest a hybrid origin of this suborder. The potential introgression scenario for Anisozygoptera is yet to be formally tested using genome-wide data.

Here we present a comprehensive analysis of transcriptomic data from 83 odonate species. First, we reconstruct a robust phylogenetic backbone using up to 4341 genetic loci for the order and discuss its evolutionary history spanning from the Carboniferous period (∼360 mya) to present day. Furthermore, in light of the “intermediate” phenotypic nature of Anisozygoptera, we investigate phylogenetic signatures of introgression within Odonata. Most notably, we identify a strong signal of deep introgression in the Anisozygoptera suborder, species of which possess traits of both main suborders, Anisoptera and Zygoptera. Although the strongest signatures of introgression are found in Anisozygoptera, we find evidence that introgression was pervasive in Odonata throughout its entire evolutionary history.

## RESULTS

### Phylogenetic Inference

We compiled transcriptomic data for 83 odonate species including 49 new transcriptomes sequenced for this study (Data S1). To assess effects of various steps of our phylogenetic pipeline on species tree inference, we examined different methods of sequence homology detection, multiple sequence alignment strategies, postprocessing filtering procedures and tree estimation methods. Specifically, three types of homologous loci (gene clusters) were used to develop our supermatrices, namely 1603 conserved single-copy orthologs (CO), 1643 all single-copy orthologs (AO) and 4341 paralogy-parsed orthologs (PO) with ≥ 42 (∼50%) species present (for more details, see Material and Methods). To date, our data represents the most comprehensive resource available for Odonata in terms of gene sampling. Each gene cluster was aligned, trimmed and concatenated resulting in five main supermatrices, CO (DNA/AA), AO (AA) and PO (DNA/AA), which included 2,167,861 DNA (682,327 amino acid (AA) sites), 882,417 AA sites, 6,202,646 DNA (1,605,370 AA) sites, respectively. Thus, the largest alignment that we used to infer the odonate phylogeny consists of 4341 loci concatenated into a supermatrix with >6 million nucleotide sites. All supermatrices are summarized in Data S2; the inferred odonate relationships are shown in Figure S1A whereas topologies of all inferred phylogenies are plotted in Figure S1B and topologies of 1603 CO gene trees are shown in Figure S1C. Additionally, we performed nodal dating of the inferred phylogeny using 20 fossil calibration points (Data S3).

The inferred ML phylogenetic tree of Odonata using DNA supermatrix of 1603 BUSCO loci (Figure 1) was used as a primary phylogenetic hypothesis throughout this study as it agrees with the majority of relationships inferred by other methods (Figures S1A, S1B). Divergence of Zygoptera and Epiprocta (Anisozygoptera+Anisoptera) from the Most Recent Common Ancestor (MRCA) occurred in the Middle Triassic ∼242 mya (Figure 1), which is in line with recent estimates [15, 36]. Comprehensive phylogenetic coestimation of subordinal relationships within Odonata showed that the suborders were well supported (Figure S1A), as they were consistently recovered as monophyletic clades in all analyses. In several previous studies, paraphyletic relationships of Zygoptera had been proposed based on wing vein characters derived from fossil odonatoids and extant Odonata [37], analysis of 12S [38], analysis of 18S, 28S, Histone 3 (H3) and morphological data [39] and analysis of 16S and 28S data [40]. In most of these studies, Lestidae was inferred to be sister to Anisoptera. Functional morphology comparisons of flight systems, secondary male genitalia and ovipositors also supported a non-monophyletic Zygoptera with uncertain phylogenetic placement of multiple groups [41]. Nevertheless, the relationships inferred from these previous datasets seem to be highly unlikely due to apparent morphological differentiation (e.g., eye spacing, body robustness, wing shape) between the suborders and support for monophyletic Anisoptera and Zygoptera from more recent morphological [34], molecular [15, 18, 42, 43] and combined studies using both data types [44]. Our analyses recover Zygoptera as monophyletic consistently (Figure S1A).

**Figure. 1.**
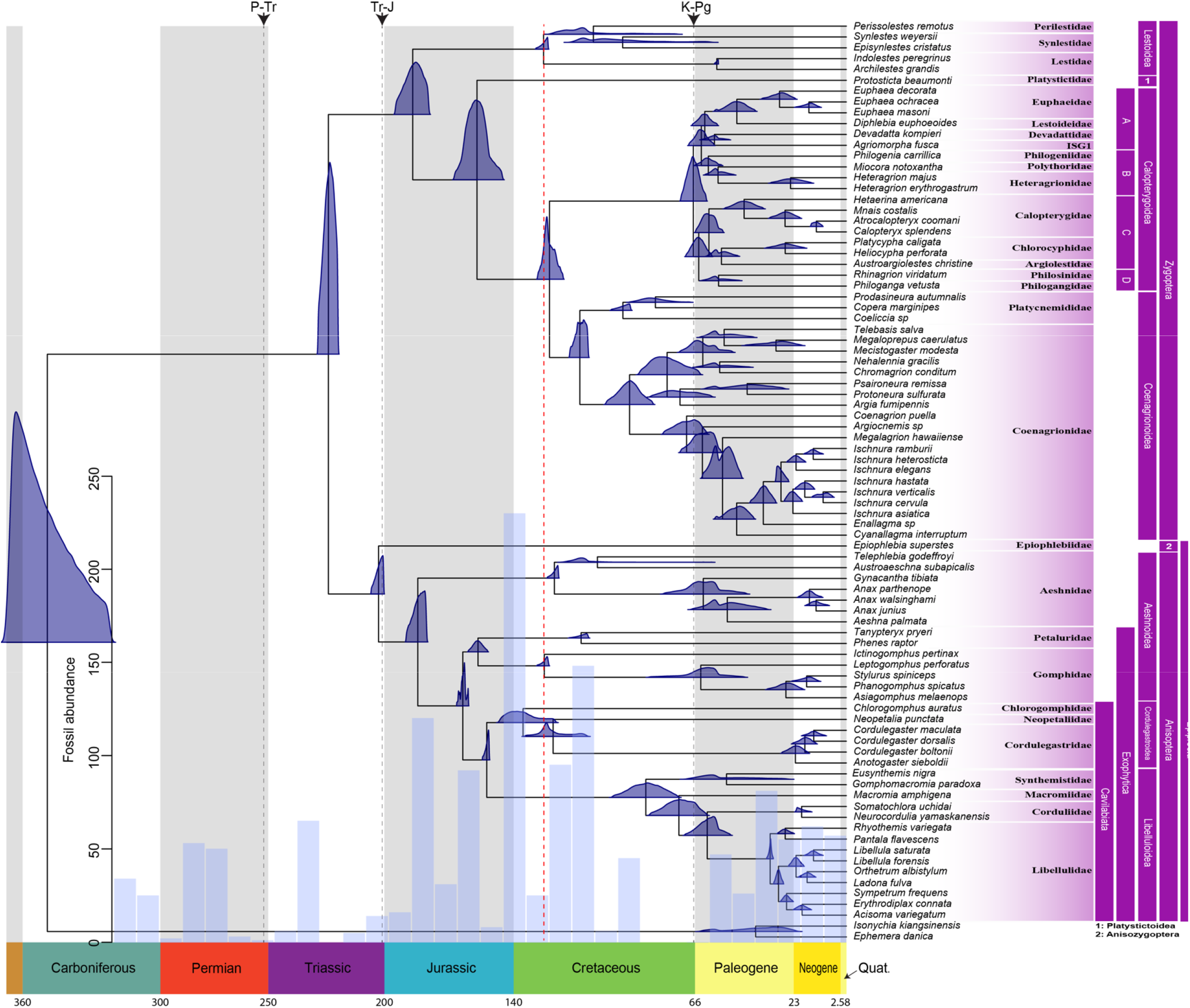
Evolutionary History of Odonata. Fossil calibrated ML phylogenetic tree of Odonata using a DNA supermatrix consisting of 1603 BUSCO genes with a total of 2,167,861 aligned sites. The blue densities at each node represent posterior distributions of ages estimated in MCMCTree using 20 fossil calibration points. The red dashed vertical line indicates the beginning of establishment for major odonate lineages originating in and spanning the Cretaceous. The histogram (blue bars) represents temporal distribution of fossil Odonatoptera samples. The black dashed vertical lines mark major extinction events, namely Permian-Triassic (P-Tr, ∼ 251 mya), Triassic-Jurassic (Tr-J, ∼ 201.3 mya) and Cretaceous-Paleogene (K-Pg, ∼66 mya)

Divergence time estimates suggest a TMRCA of Anisoptera and Anisozygoptera (we occasionally refer these two suborders as “Epiprocta”) in the Late Triassic (∼205 mya; Figure 1). Epiprocta as well as Anisoptera were consistent with more recent studies and recovered as monophyletic with very high support. We also note here that our divergence time estimates of Anisoptera tend to be younger than those found in ref. [45]. The fossil-calibration approach based on penalized likelihood has been shown to overestimate true nodal age [46] preventing direct comparison between our dates and those estimated in ref. [45].

The phylogenetic position of Gomphidae and Petaluridae, both with respect to each other and the remaining anisopteran families, has long been difficult to resolve. Several phylogenetic hypothesis have been proposed in the literature based on molecular and morphological data regarding the placement of Gomphidae as sister to the remaining Anisoptera [47] or to Libelluloidea [48]. Petaluridae has exhibited stochastic relationships with different members of Anisoptera, including sister to Gomphidae [48], sister to Libelluloidea [43], sister to Chlorogomphidae+Cordulegasteridae [44] and sister to all other Anisoptera [37, 49]. The most recent analyses of the major anisopteran lineages using several molecular markers [14] suggest Gomphidae and Petaluridae as a monophyletic group, but without strong support. Here the majority of our supermatrix analyses (Figure S1A) strongly support a sister relationship between the two families, and in our phylogeny (Figure 1) they split from the MRCA ∼165 mya in the Middle Jurassic (Figure 1); however, almost all the coalescent-based species trees reject such a relationship with high confidence (Figure S1A). In the presence of incomplete lineage sorting, concatenation methods can be statistically inconsistent [50] leading to an erroneous species tree topology with unreasonably high support [51]. Thus, inconsistency in the recovery of a sister group relationship between Gomphidae and Petaluridae can be explained by elevated levels of incomplete lineage sorting between the families and/or possible introgression events [52] (see below).

New zygopteran lineages originated in the Early Jurassic ∼191 mya with the early split of Lestoidea and the remaining Zygoptera (Figure 1). Subsequent occurrence of two large zygopteran groups, Calopterygoidea and Coenagrionoidea, was estimated within the Cretaceous (∼145-66 mya) and culminated with the rapid radiation of the majority of extant lineages in the Paleogene (∼66-23 mya). Our calibrated divergences generally agree with estimates in [15]. However, any further comparisons are precluded by the lack of comprehensive divergence time estimation for Odonata in the literature. The backbone of the crown group Calopterygoidea that branched off from Coenagrionoidea ∼129 mya in the Early Cretaceous was well supported as monophyletic in most of our inferred phylogenies (Figures 1, S1A). Previous analyses struggled to provide convincing support for the monophyly of the superfamily [13, 43, 44], whereas only 11 out of 48 phylogenetic reconstructions rejected Calopterygoidea (Figure S1A).

We used quartet sampling [53] to provide additional information about nodal support and investigate biological explanations for alternative evolutionary histories that received some support. We found that for most odonate key radiations, the majority of quartets (i.e. Frequency > 0.5) support the proposed phylogenetic hypothesis with Quartet Concordance (QC) scores > 0 (Figure S2) across all estimated putative species trees (Figure S1B). The few exceptions consist of the Gomphidae+Petaluridae split and the A+B split, where we have Frequency < 0.5 and QC < 0, which suggests that alternative relationships are possible. Quartet Differential (QD), inspired by *f* and *D* statistics for introgression, provides an indicator of how much the proportions of discordant quartets are skewed (i.e. whether one of the two discordant relationships is more common than the other), suggestive of introgression and/or introgression or substitution rate heterogeneity [53]. Interestingly, we identified skewness (i.e. QD < 0.5) for almost every major radiation (Figure S2), which suggests that alternative relationships can be a result of underlying biological processes (e.g. introgression) rather than ILS alone [53]; however, this score may not be highly informative if the majority of quartets agree with the focal topology (i.e Frequency > 0.5 and QC > 0). For both the Gomphidae+Petaluridae and the A+B splits, we have Frequency > 0.5, QC < 0 and QD < 0.5, implying that alternative phylogenetic relationships are plausibly not only due to ILS but also possible ancestral introgression.

### Major Trends in Evolutionary History of Odonata

Investigation of diversification rates in Odonata highlighted two major trends correlated with two mass extinction events in the Permian-Triassic (P-Tr) ∼251 mya and Cretaceous-Paleogene (K-Pg) ∼66 mya. First, it appears likely that the ancient diversity of Odonata was largely eliminated after the P-Tr mass extinction event (see the temporal distribution of fossil samples in Figure 1) as was also the case for multiple insect lineages [54]. According to the fossil record at least two major odonatoid lineages went extinct (Protodonata and Protanisoptera [55]) and likely many genera from other lineages as well (e.g., Kargalotypidae from Triadophlebimorpha [56]). The establishment of major odonate lineages was observed during the Cretaceous starting ∼135 mya (Figure 1, red line). This coincided with the radiation of angiosperm plants that, in turn, triggered the formation of herbivorous insect lineages [36]. Odonates are exclusively carnivorous insects and their diversification was likely driven by the aforementioned sequence of events. Interestingly, molecular adaptations in the odonate visual systems are coupled with their diversification during the Cretaceous as well [18].

### Overview of introgression hypotheses tested

The scope of introgression within Odonata remains largely unknown, where previous studies looked for its patterns only within certain species relying on inference from a limited number of genetic loci. Thus, we searched for signatures of introgression using genome-scale datasets between lineages at several different taxonomic levels: between different suborders, between superfamilies and within superfamilies. We used six different methods to test for introgression within Odonata, as exemplified for the Anisozygoptera suborder in Figure 2. Specifically, we searched for signatures of ancestral inter-superfamilial introgression within the Zygoptera and Anisoptera suborders. Also, we tested the hypothesis of inter-subordinal gene flow between Anisozygoptera and Zygoptera. Finally, we tested introgression within superfamilies of Zygoptera and Anisoptera that included several species (Lestoidea, Calopterygoidea, Coenagrionoidea, Aeshnoidea (Aeshnidae), Aeshnoidea (Gomphidae+Petaluridae), Cordulegastroidea and Libelluloidea; Figure 1). The introgression results for the entire Odonata order either comprise of all tests performed within the entire phylogeny (HyDe) or of a union of tests performed within Anisoptera, Zygoptera and between Anisozygoptera-Zygoptera (*D*_FOIL_, QuiBL and *χ*^2^ count test/Branch Length Test (BLT)).

**Figure 2.**
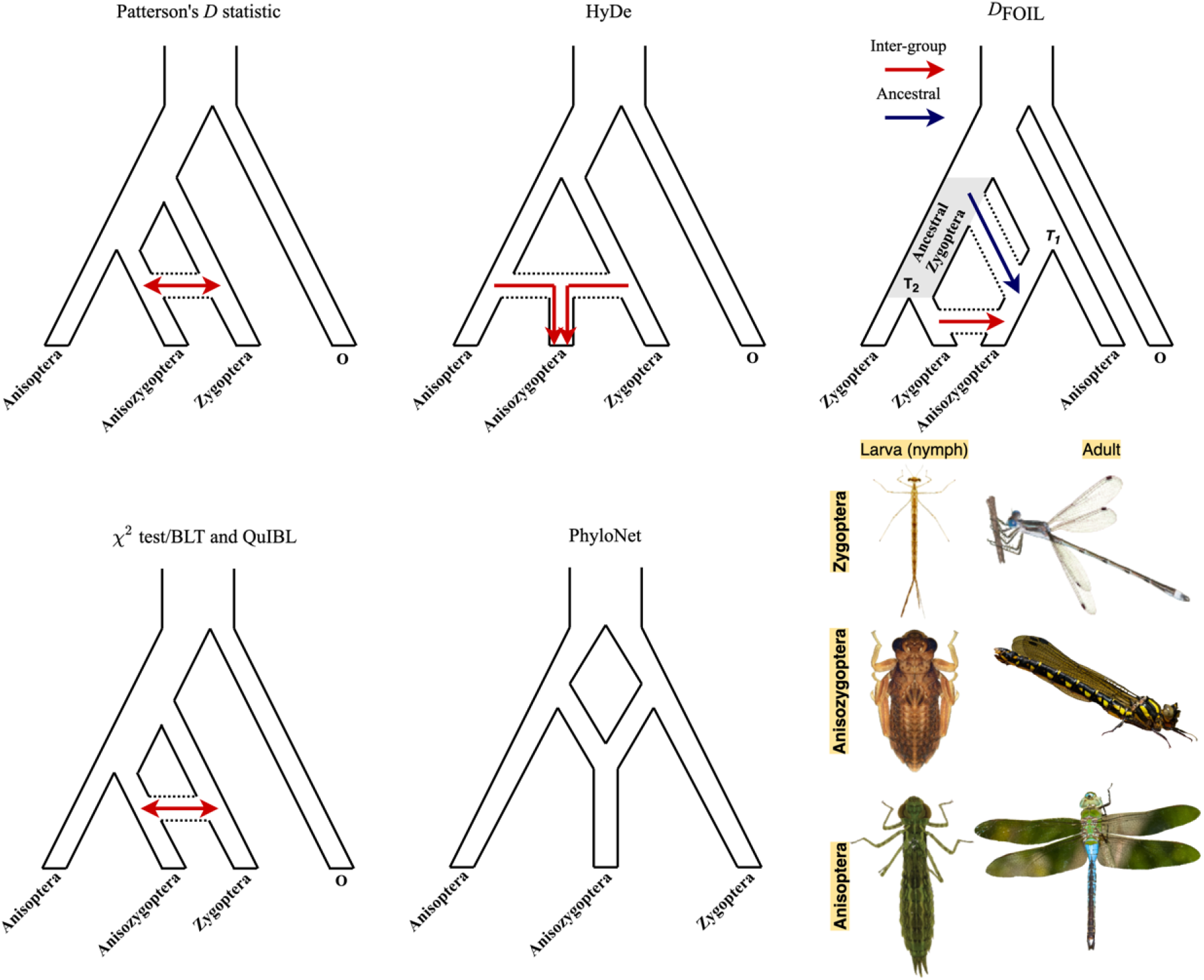
Detection of Introgression/Hybridization Trajectories by Different Methods in Anisozygoptera. Three site pattern-based (*D* statistic, HyDe and *D*_FOIL_), gene tree count/branch length-based (*χ*^2^ test/BLT and QuIBL) and network ML inference PhyloNet) methods were used to test for introgression/hybridization. Arrows denote introgression. The figure panel represents larval and adult stages for three Odonata suborders. Species from top to bottom: *Lestes australis, Epiophlebia superstes* and *Anax junius*. Image credit: *Epiophlebia superstes* adult by Christian Dutto Engelmann; *Lestes australis* and *Anax junius* adults by John Abbott.

### Site-pattern-based methods strongly suggest multiple instances of introgression within Odonata

Initially, we tested the above hypotheses of introgression in quartet topologies (Figure S3) within Odonata using two site-pattern-based methods: the ABBA-BABA test [57] and HyDe [58]. The ABBA-BABA test and HyDe rely on computation of *D* and *γ* statistics, respectively, where their significant deviation from 0 may indicate the presence of introgression between the tested pair of taxa. Additionally, estimated *γ* and (1-*γ*) of HyDe’s hybrid speciation model [58] corresponds to the parental fractions in a putatively hybrid genome. Note that HyDe’s hybrid speciation model is appropriate for detecting introgression with sufficient statistical power and can produce reasonable estimates of *γ* [59, 60]. The analysis of the ABBA-BABA test results revealed possible gene flow events throughout the entire evolutionary history of Odonata (Figure S4 A, Data S4, SI Results). We also highlight the positive relationship between the values of *D* and *γ* statistics (Spearman’s rank correlation test, ρ = 0.308, *P* = 0), demonstrating their broad concordance in identifying signatures of introgression (Figure S4 B).

There are a variety of other test statistics that have been developed to detect introgression (e.g., [61-64]). Because, like *D* and *γ*, many of these statistics are computed from different invariants, we attempted to visualize the relationships between all 15 site patterns computed by HyDe using *t*-distributed stochastic neighbor embedding (tSNE; [65]) for dimensionality reduction, along with the corresponding values of *D* and *γ* (Figures S4 C and D, respectively). We found that the clustering of quartets with significant introgression according to a two-dimensional representation of their site patterns may suggest the presence of additional site-pattern signatures of introgression (Figure S4 C-D).

In order to assess the extent of preservation of ancestrally introgressed genetic material within contemporary taxa, we compared inferred average values of significant *γ* statistic from Odonata with the averages derived from different intra- and inter-superfamilial taxonomic levels (Figure 3A). We found significantly higher values of *γ* for several intra- and inter-superfamilial comparisons including those that involve Anisozygoptera (Wilcoxon rank-sum test [WRST], all *P* < 0.05, Figure 3A, Data S4, SI Results). Additionally, Anisozygoptera exhibits the largest average *γ* (0.27) across all the inter-superfamilial comparisons (Figure 3A, Data S4). We also found an excess (Fisher exact test [FET], all *P* < 0.05, Data S1) of significant triplets that support introgression (Figure 3B) based on both the ABBA-BABA and HyDe hybrid speciation model (Figure S3) tests for Anisozygoptera, Aeshnoidea (Aeshnidae), Lestoidea and Calopterygoidea (between and within). Additionally, HyDe analysis yielded highly similar results for *D* and *γ* using only the 1^st^ and 2^nd^ codon positions (Figure S5) suggesting that the potential saturation effect in 3^rd^ codon position had a minor effect on introgression inference.

**Figure 3.**
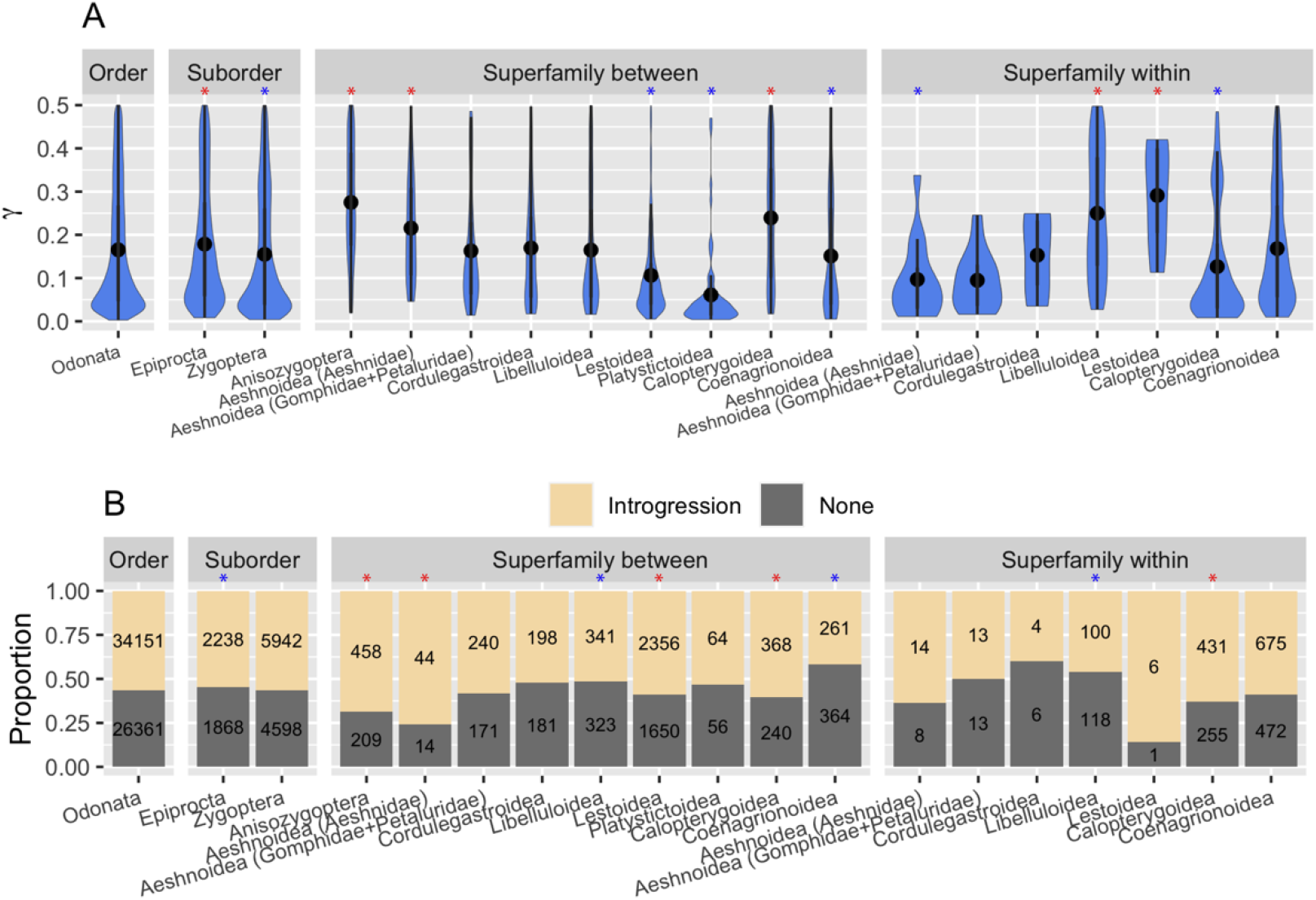
Distributions of HyDe γ and Quartets Fractions Supporting Introgression Across Odonate Taxonomic Levels. (A) Distribution of significant (Bonferroni corrected *P* < 10^−6^) γ values for each quartet estimated by HyDe. In general, γ values that are not significantly different from 0 denote no relation of a putative hybrid species to either of the parental species P_1_ (1-γ) or P_2_ (γ) in a quartet. Asterisks indicate significantly greater (red) or lower (blue) γ averages of various tested cases compared to γ average of the entire order. (B) Proportions of quartets that support introgression based on simultaneous significance of *D* statistics and γ. Asterisks indicate significantly greater (red) and smaller (blue) fraction of quartets that support introgression compared to the entire order.

Further, we tested introgression within Odonata using an alternative invariant-based method, *D*_FOIL_ (Figure S6 A, SI Results), which is capable of detecting ancestral as well as inter-group introgression and inferring its polarization (see Materials and Methods). Specifically, we observed a highly skewed distribution of *D*_FOIL_ statistics for Anisozygoptera (Figure S6 B, SI Results) that may suggest specific directionality of introgression: all 61 (1157 out of 1158 without FDR correction) quintets with positive evidence for ancestral introgression suggest that Anisozygoptera is the recipient lineage whereas Zygoptera is the donor (Figure 2, red and blue arrows). Additionally, 1030 out of 1091 (4473 out of 4738 without FDR correction) evaluated quintets with significant inter-group introgression are indicative of one-way introgression from Zygoptera lineages to Anisozygoptera as well, whereas the directionality of remaining 59 (254 without FDR correction) quintets support Anisozygoptera as a donor and Zygoptera as a recipient and only two (11 without FDR correction) show introgression between Zygoptera and Anisoptera.

### Signatures of introgression revealed by phylogenetic gene tree-based discordance methods

Besides specific phylogenetic invariants, introgression will also generate certain patterns of gene tree-species tree discordance and reduce the genetic divergence between introgressing taxa, which is reflected in gene tree branch lengths. Thus, the footprints of introgression can be detected using phylogenetic discordance methods. First, for a triplet of species, under ILS alone one would expect equal proportions of gene tree topologies supporting the two topologies disagreeing with the species tree, and any imbalance may suggest introgression (Figure S7). Thus, deviation from equal frequencies of gene tree counts among discordant gene trees can be assessed using a *χ*^2^ test, similar to the method proposed in [66, 67]. One would expect the average distance between putatively introgressing taxa in discordant trees to be significantly smaller than the distances derived from the concordant as well as alternative discordant triplets (see Materials and Methods and Figure S7). We used both the *χ*^2^ test of discordant gene tree counts and a test based on the distribution of branch lengths for concordant and discordant gene tree triplets (BLT) to identify introgression within Odonata testing different scenarios (Figure 4A). With this combination of methods, we identified a significant fraction of triplets that support ancestral introgression for scenario 1 that involves inter-subordinal gene flow between Anisozygoptera and Zygoptera as well as scenarios 2 through 4, which correspond to inter-superfamilial instances of introgression within Epiprocta (Figure 4B). Within superfamilies we found signatures of introgression for Calopterygoidea and Libelluloidea (Figure 4B). For scenario 1 (introgression between Zygoptera and Anisozygoptera), examination of the genetic divergence distributions (Figure 4C) for concordant and discordant triplets showed that discordant triplets that may have resulted from gene flow between Anisozygoptera and Zygoptera (the topology labeled “discord2”), have markedly smaller average divergence between these two taxa, as expected in the presence of introgression. Similarly, based on the distribution of mean divergence between putatively introgressing taxa (Figure S8) as well as fraction of significant triplets (Figure 4B) gene flow is supported for scenarios 2, 3 and 4.

**Figure 4.**
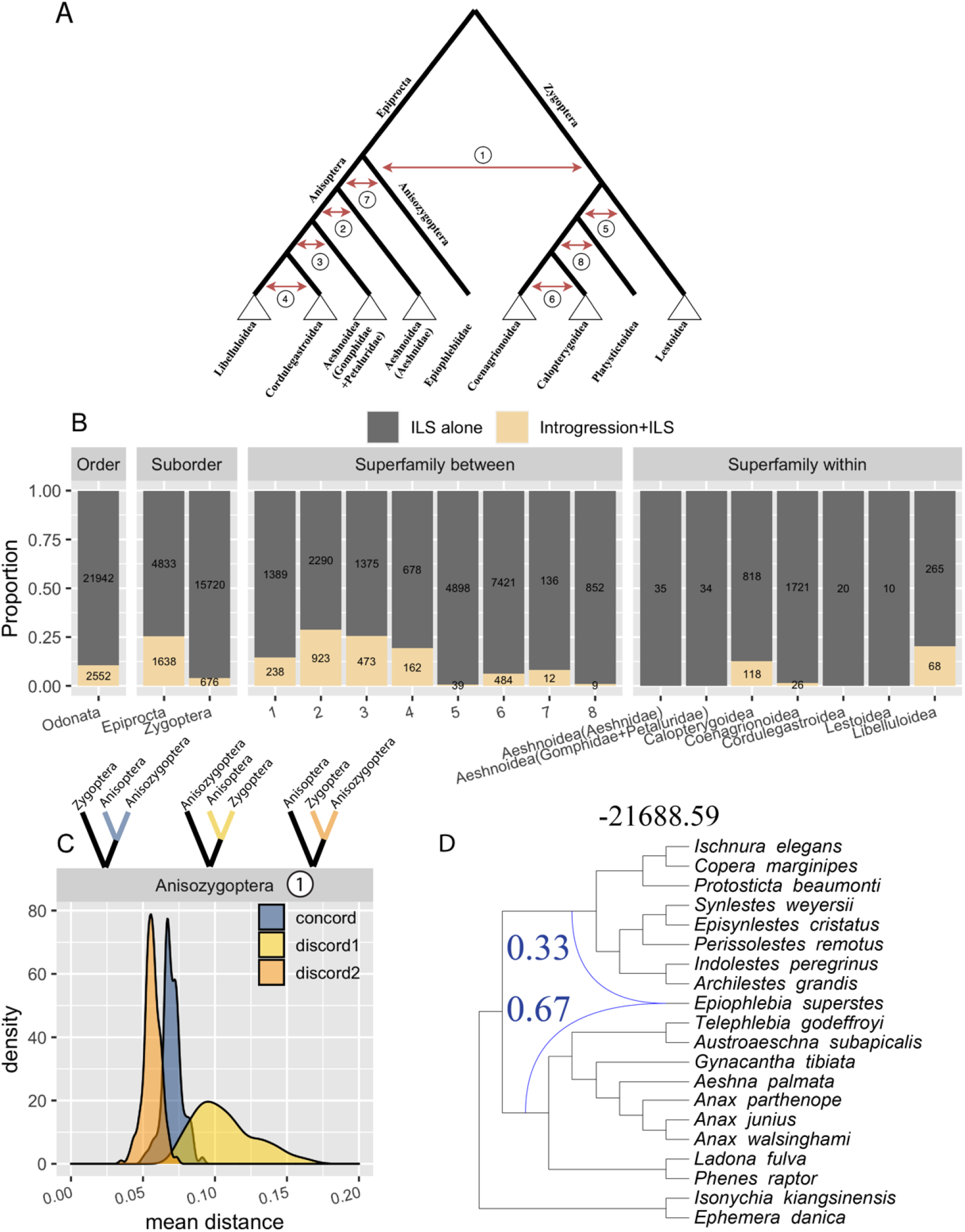
Results of the *χ*^2^ Count-Branch Length Test (BLT) for Odonate Taxonomic Levels and PhyloNet result for Anisozygoptera. (A) Scenarios of deep (numbered red arrows) and intra-superfamilial (white triangles) introgression in Odonata tested using *χ*^***2***^ test and BLT. Red arrows mark the location of ancestral introgression events that were tested between lineages (e.g. for scenario 8, we tested whether contemporary species of Platystictoidea shared introgressed genetic material with either Coenagrionoidea or Calopterygoidea). (B) Classification of triplets based on the *X*^2^ test and BLT results. Introgression+ILS cases are those significant according to both the *χ*^2^ test and BLT (FDR corrected *P <* 0.05); all the remaining cases were those where any discordance was inferred to be due to ILS alone. (C) Normalized genetic divergence between sister taxa (shown in triplets above the panel) averaged across all BUSCO gene trees supporting each topology involving three lineages representing Anisoptera, Anisozygoptera, and Zygoptera. (D) Phylogenetic network estimated from a set of ML gene trees using maximum likelihood. *Epiophlebia superstes*, the sole representative of Anisozygoptera in our study, was specified to be involved in a reticulation with two other (unspecified) lineages. Blue lines indicate the reticulation event and are labeled with PhyloNet’s estimate of γ. The number above the network indicates the log-likelihood score.

An alternative gene tree branch length-based approach, QuIBL [68], also detected multiple instances of introgression within the entire order (Figure S9). We note that particularly for introgression involving Anisozygoptera we observed a larger fraction of triplets suggestive of introgression than for any other scenario tested.

### Phylogenetic network analyses support Anisozygoptera-Zygoptera introgression

As an alternative approach to identify the lineage experiencing gene flow with Anisozygoptera, we performed phylogenetic network inference in PhyloNet [69, 70]. For this analysis we specified that Anisozygoptera was involved in a reticulation event—the only such event occurring on the tree—and inferred for the other two nodes that were most likely to be involved in the reticulation as well as the value of *γ*, the fraction of Anisozygoptera’s genetic material derived from this reticulation. The full maximum likelihood (Figure 4D) and maximum pseudolikelihood (Figure S10) approaches recovered topologically similar networks with comparable values of *γ*. Pseudolikelihood analysis suggests a reticulation event between Aeshnidae (a family within Anisoptera) and Zygoptera with 38% of genetic material coming from Zygoptera (Figure S10), whereas full likelihood infers reticulation between Anisoptera and Zygoptera suborders 33% (Figure 4D) of genetic material from Zygoptera. Overall, we note that the full likelihood approach returned networks with higher log-likelihood scores. This observation is most likely due to the fact that we performed full likelihood analysis in an exhaustive manner (Data S6) testing every possible reticulation event within a focal clade topology congruent with the species tree (Figure 1), whereas pseudolikelihood analysis used the hill-climbing algorithm to search the full network space but is not guaranteed to retrieve the most optimal solution. Moreover, differences in objective function within likelihood and pseudolikelihood frameworks could also lead to distinct network topology and *γ* estimate.

### Methodological consensus corroborates strong evidence of introgression

We examined the agreement across site- and gene tree-based methods for detecting introgression (i.e. Hyde/*D*, and *D*_FOIL_ and the *χ*^2^ test/BLT) and summarize the results in (Figure S11).

Specifically, we compared sets of unique introgressing species pairs that were identified with significant pattern of introgression. Overall, all methods individually were able to identify abundant introgression within the entire Odonata order with strong agreement identified by the significance of overlap using an exact test of multi-set interactions [71]. We found strong overlap across methods in their support of introgression within Epiprocta (scenarios 2 and 3), gene flow involving Anisozygoptera (scenario 1), and intra-superfamilial introgression within Libelluloidea. Within Zygoptera our methods showed strong overlap in identifying signatures of introgression for Calopterygoidea only. Notably, our *χ*^2^ test /BLT produced very few predictions that were not in agreement with Hyde, *D* and/or *D*_FOIL_ across different comparisons.

## DISCUSSION

### Resolving main radiations on Odonata phylogeny

Several competing hypotheses of evolutionary relationships within Odonata have been proposed by multiple authors regarding various taxonomic levels (Figure S12A) [13, 14, 18, 37, 38, 40, 43, 44, 47, 48, 72, 73]. Our phylogenetic analyses recovered Epiprocta (Anisoptera+Anisozygoptera) and Zygoptera as monophyletic with high support (Figure S1A) agreeing with other recent estimates [18, 43, 44, 72]. Our estimated superfamilial relationships within Zygoptera support hypothesis of Dijkstra, et al. [13] recovering monophyly of Lestoidea, Platystictoidea, Coenagrionoidea and Calopterygoidea (Figure S12A) with high support (Figure S1A). Inferred higher-level phylogenetic classification of anisopteran families was highly congruent with [14] and well-supported (Figure S1A) with the exception of Gomphidae and Petaluridae radiations. The phylogenetic position of Gomphidae and Petaluridae, both with respect to each other and the remaining anisopteran families, has long been difficult to resolve (Figure S12C). The most recent analyses of major anisopteran lineages, using several molecular markers [14], suggest Gomphidae and Petaluridae as a monophyletic group, but without strong branch support. Here the majority of the supermatrix analyses strongly support a sister relationship between the two families (Figure S1A); however, almost all the ASTRAL species tree analyses reject such a relationship with high confidence (Figure S1A). In the presence of incomplete lineage sorting concatenation methods can be statistically inconsistent [50] leading to an erroneous species tree topology with unreasonably high support [51]. Thus, inconsistency in the recovery of a sister group relationship between Gomphidae and Petaluridae could be a result of elevated levels of incomplete lineage sorting between the families.

### Widespread introgression within Odonata

Introgression among Odonata has been previously thought to be uncommon [74, 75] as a result of probable reproductive isolation mechanisms such as ethological barriers, phenotypic divergence (e.g., variable morphology of genitalia [76, 77]) and habitat and temporal isolation. Most of the introgression and hybridization research in Odonata has been done within the zygopteran Coenagrionidae family and especially between populations of genus *Ishnura* focusing on mechanisms of reproductive isolation (e.g. [25, 78]). These studies established “hybridization thresholds” for these closely related species and showed positive correlation between isolation and genetic divergence. Furthermore, rapid karyotype evolution can also contribute to post-mating isolation [79]; however, recent cytogenetic studies across Odonata phylogeny indicate very stable chromosome number with most prevalent karyotype of 2n = 25 [80]. Additionally, in recent years there has been a growing body of evidence for introgression throughout evolutionary histories of various groups. This includes more recent hybridization events observed in *Heliconius* butterflies [68], cats [81], cichlid fishes [82] as well as deeper ancestral introgression in primates [83], *Drosophila* [84] and vascular plants [53].

On the shallow evolutionary timescales, the rate of gene flow and maintenance of introgressed variation can be approximated by the fraction of introgressed material (i.e. by *γ* in our case) in a focal genome [85]. However, with the increased divergence time the fixation or loss of the ancestrally introduced genetic material will primarily depend on the scope of selection [6] via its adaptive value [7, 8]. Here, our taxon sampling allowed to test ultra-deep ancestral introgression scenarios where the MRCA of tested taxa can be traced as far back as to the Triassic period (∼251 mya to ∼201 mya). We argue that for the cases where we infer that a large amount of introgressed genetic material was preserved (e.g. >25% in Anisozygoptera) the ancestral effective migration rates [85] were high (similar conclusions can be made for Aeshnoidea (Aeshnidae), Calopterygoidea and within Libelluloidea and Lestoidea; Figure 3A); whereas for the remaining taxa with lower or non-significant deviations of *γ* (Figure 3A) the rates of introgression may have been substantially lower or its signatures were purged from the genome by selection.

We found compelling evidence for patterns of ancestral introgression (Figure S11) among distantly related odonate taxa. Our conservative analysis of agreement between different introgression tests shows presence of gene flow within the entire order (Figure S11). This observation may be explained by reduced sexual selection pressures in the early stages of odonate evolution that may have inhibited rapid genital divergence [86], which is probably a primary source of reproductive isolation in Odonata [87].

### Ancestral introgression as an evolutionary driver in Anisozygoptera

Perhaps the most compelling evidence that we uncovered was for deep ancestral introgression between the Zygoptera and Anisozygoptera suborders, which based on the fossil record became genetically isolated after the Lower Jurassic [88]. Species of Anisozygoptera exhibit anatomical characteristics of both Anisoptera and Zygoptera suborders. Some general features of Anisozygoptera that relate them to Zygoptera include dorsal folding of wings during perching in adults, characteristic anatomy of proventriculus (a part of digestive system that is responsible for grinding of food particles) and absence of specific muscle groups in the larval rectal gills; whereas abdominal tergite shape, rear wing geometry and larval structures are similar to Anisoptera [89]. More recent studies also revealed that Anisozygoptera ovipositor morphology shares similarity with Zygoptera [90]; muscle composition of the head resemble characteristics of both Anisoptera and Zygoptera [91]; thoracic musculature of Anisozygoptera larva exhibit similarity between Anisoptera and Zygoptera [34]. Thus, Anisozygoptera represent a morphological and behavioral “intermediate” [92], which is supported by our findings where we consistently recovered introgression between Zygoptera and Anisozygoptera (Figures 2-4). Moreover, according to the *D*_FOIL_ method the Anisozygoptera was inferred to be a recipient taxon from a zygopteran donor, though we cannot rule out a lesser degree of gene flow in the opposite direction as well. Strikingly, the average HyDe probability parameter *γ ∼* 0.27 (Figure 3A) inferred from multiple quartets is very similar to PhyloNet’s inheritance probability *γ ∼* 0.33. This may suggest that a sizeable fraction of the Anisozygoptera genome descends from zygopteran lineages. Taken together, these observations strongly suggest xenoplasious origin [93] of Anisozygopteran traits (i.e. traits introduced to a recipient taxon via introgression) that are shared with Zygoptera. However, we do not reject the possibility that some trait hemiplasy may have resulted from ILS [94]. Based on the gathered evidence for introgression, we suggest that ancestral lineages that gave rise to modern day Anisozygoptera and Zygoptera taxa experienced introgression in their past evolutionary history.

According to the hybrid swarm hypothesis [95], introgression can trigger a rapid cladogenesis resulting in the establishment of ecologically different species with niche-specific adaptive phenotypes. Some of the notable examples can include adaptive radiations caused by ancestral introgression in cichlid fishes [96], and intricate inter-specific introgression patterns in big cats that could be linked to rapid diversification of modern-day species in that lineage [97]. Interestingly, the large fraction of introgressed material estimated within the Anisozygopteran lineage does not appear to have facilitated any adaptive radiation. This proposition is supported by an extremely low species diversification within the suborder (just three extant species) and also by the lack of fossil record for Anisozygoptera.

## Conclusions

Our findings provide the first insight into global patterns of introgression that were pervasive throughout the evolutionary history of Odonata. Further work based on full genome sequencing data, including non-coding regions, and denser taxon sampling will be necessary to more accurately estimate the fraction of the introgressed genome in various species. We also stress the importance of further method development to allow for more accurate and scalable inference of phylogenetic networks and enable researchers to pin-point instances of inter-taxonomic introgression on the phylogeny.

## ACKNOWLEDGMENTS

We thank Gavin Martin, Nathan Lord and Camilla Sharkey for the generation of sequence data. We also thank the DNA Sequencing Center and Fulton Supercomputer Lab (BYU) for assistance. We also thank Aaron Comeault for his invaluable comments on the manuscript. This work was supported by the NSF grant DEB-1265714 to SMB and NIH grants R00HG008696 and R35GM138286 to DRS. CS was funded by grant ANR-19-CE45-0012 from French Agence Nationale de la Recherche.

## AUTHOR CONTRIBUTIONS

AS, CS, DRS and SMB conceived the research. AS, CS, MSF and PB analyzed the data. MC, KAC, MFW and DRS supervised the project. AS, CS, DRS and SMB wrote the manuscript.

## MATERIALS AND METHODS

### Taxon sampling and RNA-seq

In this study, we used distinct 85 species (83 ingroup and 2 outgroup taxa). 35 RNA-seq libraries were obtained from NCBI (see Data S1). The remaining 58 libraries were sequenced in Bybee Lab (some species have several RNA-seq libraries; Data S1). Total RNA was extracted for each taxon from eye tissue using NucleoSpin columns (Clontech) and reverse-transcribed into cDNA libraries using the Illumina TruSeq RNA v2 sample preparation kit that both generates and amplifies full-length cDNAs. Prepped mRNA libraries with insert size of ∼200bp were multiplexed and sequenced on an Illumina HiSeq 2000 producing 101-bp paired-end reads by the Microarray and Genomic Analysis Core Facility at the Huntsman Cancer Institute at the University of Utah, Salt Lake City, UT, USA. Quality scores, tissue type and other information about RNA-seq libraries are summarized in Data S1.

### Transcriptome assembly and CDS prediction

RNA-seq libraries were trimmed and de novo assembled using Trinity [98, 99] with default parameters. Then only the longest isoform was selected from each gene for downstream analyses using Trinity utility script. In order to identify potentially coding regions within the transcriptomes, we used TransDecoder with default parameters specifying to predict only the single best ORF. Each predicted proteome was screened for contamination using DIMOND BLASTP [100] with an E-value cutoff of 10^−10^ against custom protein database. Non-arthropod hits were discarded from proteomes (amino acid, AA sequences) and corresponding CDSs. To mitigate redundancy in proteomes and CDSs, we used CD-HIT [101] with the identity threshold of 0.99. Such a conservative threshold was used to prevent exclusion of true paralogous sequences; thus, reducing possible false positive detection of 1:1 orthologs during homology searches.

### Homology assessment

In the present study three types of homologous loci (gene clusters) namely conserved single-copy orthologs (CO), all single-copy orthologs (AO) and paralogy-parsed orthologs (PO) identified by BUSCO v1.22 [102], OrthoMCL [103] and Yang’s [104] pipelines respectively were used in phylogenetic inference.

BUSCO arthropod Hidden Markov Model Profiles of 2675 single-copy orthologs were used to find significant COs matches within CDS datasets by HMMER’s *hmmersearch* v3 [105] with group-specific expected bit-score cutoffs. BUSCO classifies loci into complete [duplicated] and fragmented. Thus, only complete single-copy loci were extracted from CDS datasets and corresponding AA sequences for further phylogenetic analyses. Since loci were identified as true orthologs if they score above expected bit-score and complete if their lengths lie within ∼95% of BUSCO group mean length, many partial erroneously assembled sequences were filtered out.

OrthoMCL v2.0.9 [103] was used to compute AOs in all species using predicted AA sequences by TransDecoder. AA sequences were used in an all-vs-all BLASTP with an E-value cutoff of 10^−10^ to find putative orthologs and paralogs. The Markov Cluster algorithm (MCL) inflation point parameter was set to 2. Only 1:1 orthologs were used in further analyses. In order to exclude false-positive homology clusters identified by OrthoMCL, we applied machine learning filtering procedure [106] implemented in OGCleaner software v1.0 [107] using a metaclassifier with logistic regression.

Finally, to identify additional clusters, we used Yang’s tree-based orthology inference pipeline [104] that was specifically designed for non-model organisms using transcriptomic data. Yang’s algorithm is capable of parsing paralogous gene families into “orthology” clusters that can be used in phylogenetic analyses. It has been shown that paralogous sequences encompass useful phylogenetic information [108]. First, the Transdecoder-predicted AA sequences were trimmed using CD-HIT with the identity threshold of 0.995. Then, all-vs-all BLASTP with an E-value cutoff of 10^−5^ search was implemented. The raw BLASTP output was filtered by hit fraction of 0.4. Then, MCL clustering was performed with inflation point parameter of 2. Each cluster was aligned using iterative algorithm of PASTA [109] and then was used to infer a maximum-likelihood (ML) gene tree using IQ-TREE v1.5.2 [110] with an automatic model selection. Tree tips that were longer than relative and absolute cutoffs of 0.4 and 1 respectively were removed. Mono- and paraphyletic tips that belonged to the same species were masked as well. To increase quality of homology clusters realignment, tree inference and tip masking steps were iterated with more stringent relative and absolute masking cutoffs of 0.2 and 0.5 respectively. Finally, POs (AA sequences and corresponding CDSs) were extracted by rooted ingroups (RI) procedure using *Ephemera danica* as an outgroup (for details see [104]).

### Cluster alignment, trimming and supermatrix assembly

For most of the analyses only clusters with ≥ 42 (∼50%) species present were retained. In total, we obtained five cluster types, namely DNA (CDS) and AA COs, AA AOs and DNA and AA POs. Each cluster was aligned using PASTA [109] for the DNA and AA alignments and PRANK v150803 [111, 112] for the codon alignments and alignments with removed either 1^st^ and 2^nd^ or 3^rd^ codon positions. In order to reduce the amount of randomly aligned regions, we implemented ALISCORE v2.0 [113] trimming procedure (for PASTA alignments) followed by masking any site with ≥ 42 gap characters (for both PASTA and PRANK alignments). Also, sequence fragments with >50% gap characters were removed from clusters that were subjected to ASTRAL v4.10.12 [114] estimation since fragmentary data may have a negative effect on accuracy of gene and hence species tree inference [115]. For each of the cluster type, we assembled supermatrices from trimmed gene alignments. Additionally, completely untrimmed supermatrices were generated from DNA and AA COs with ≥ 5 species present.

### Phylogenetic tree reconstruction

Four contrasting tree building methods (ML:IQTREE, Bayesian:ExaBayes, Supertree:ASTRAL, Alignment-Free (AF): Co-phylog) were used to infer odonate phylogenetic relationships using different input data types (untrimmed and trimmed supermatrices, codon supermatrices, codon supermatrices with 1^st^ and 2^nd^ or 3^rd^ positions removed, gene trees and assembled transcriptomes). In total we performed 48 phylogenetic analyses and compared topologies to identify stable and conflicting relationships (Data S2).

We inferred phylogenetic ML trees from each supermatrix using IQTREE implementing two partitioning schemes: single partition and those identified by PartitionFinder v2.0 (three GTR models for DNA and a large array of protein models for AA) [116] with relaxed hierarchical clustering option [117]. In the first case, IQTREE was run allowing model selection and assessing nodal support with 1000 ultrafast bootstrap (UFBoot) [118] replicates. In the second case, IQTREE was run with a given PartitionFinder partition model applying gene and site resampling to minimize false-positives [119] for 1000 UFBoot replicates.

For Bayesian analyses implemented in ExaBayes [120], we used highly trimmed (retaining sites only with occupancy of ≤ 5 gap characters) and original DNA and AA CO supermatrices assuming a single partition. We initiated 4 independent runs with 4 Markov Chain Monte Carlo (MCMC) coupled chains sampling every 500^th^ iteration. Due to high computational demands of the procedure, only the GTR and JTT substitution model priors were applied to DNA and AA CO supermatrices respectively with the default topology, rate heterogeneity and branch lengths priors. However, all supported protein substitution models as a prior was specified for the trimmed AA CO supermatrix. For convergence criteria an average standard deviation of split frequencies (ASDSF) [121], a potential scale reduction factor (PSRF) [122] and an effective sample size (ESS) [123] were utilized. Values of 0% < ASDSF <1% and 1% < ASDSF < 5% indicate excellent and acceptable convergence respectively; ESS >100 and PSRF ∼ 1 represent good convergence (see ExaBayes manual, [120]).

ASTRAL analyses were conducted using two input types: (i) gene trees obtained by IQ-TREE allowing model selection for fully trimmed DNA and AA clusters and (ii) gene trees obtained from the alignment-tree coestimation process in PASTA. Nodal support was assessed by local posterior probabilities [124]. ASTRAL is a statistically consistent supertree method under the multispecies coalescent model with better accuracy than other similar approaches [114].

In addition to standard phylogenetic inferential approaches, we applied an alignment-free (AF) species tree estimation algorithm using Co-phylog [125]. Raw Transdecoder CDS outputs were used in this analysis using k-mer size of 9 as the half context length required for Co-phylog. Bootstrap replicate trees were obtained by running Co-Phylog with the same parameter settings on each subsampled with replacement CDS Transdecoder libraries and were used to assess nodal support.

### Assessment of phylogenetic support via quartet sampling

As an additional phylogenetic support, we implemented quartet sampling (QS) approach [53]. Briefly, it provides three scores for internal nodes: (i) the quartet concordance (QC) score gives an estimate of how sampled quartet topologies agree with the putative species tree; (ii) quartet differential (QD) estimates frequency skewness of the discordant quartet topologies, which can be indicative of introgression if a skewed frequency is observed and (iii) quartet informativeness (QI) quantifies how informative sampled quartets are by comparing likelihood scores of alternative quartet topologies. Finally, QS provides a quartet fidelity (QF) score for terminal nodes that measures a taxon “rogueness”. We performed QS analysis with all 48 putative species phylogenies using SuperMatrix_50BUSCO_dna_pasta_ali_trim supermatrix, specifying an IQ-TREE engine for quartet likelihood calculations with 100 replicates (i.e. number of quartet draws per focal branch).

### Fossil dating

A Bayesian algorithm of MCMCTREE [126] with approximate likelihood computation was implemented to estimate divergence times within Odonata using 20 crown group fossil constraints with corresponding prior distributions (Data S3). First, we estimated branch length by ML and then the gradient and Hessian matrix around these ML estimates in MCMCTREE using SuperMatrix_50BUSCO_dna_pasta_ali_trim supermatrix. Second, we used the gradient and Hessian matrix which constructs an approximate likelihood function by Taylor expansion [127] to perform fossil calibration in MCMC framework under the uncorrelated clock model. For this step we specified GTR+Γ substitution model with 4 gamma categories; birth, death and sampling parameters of 1, 0.5 and 0.01, respectively. To ensure convergence, the analysis was run independently 10 times for 10^6^ generations, logging every 50th generation and then removing 10% as burn-in. Visualization of the calibrated tree was performed in R using MCMCtreeR package [128].

### Analyses of introgression

In order to address the scope of possible reticulate evolution across odonate phylogeny, we used various methods of introgression detection such as HyDe/*D* [59], *D*_FOIL_ [129], *X*^2^ goodness-of-fit test, QuIBL [68] and PhyloNet [69, 70].

The HyDe framework allows detection of hybridization events which relies on quantification of phylogenetic site patterns (invariants). HyDe estimates whether a putative hybrid population (or taxon) H is sister to either population P1 with probability γ or to P2 with probability 1-γ in a 4-taxon (quartet) tree ((P1,H,P2),O), where O denotes an outgroup. Then, it conducts a formal statistical test of H_0_: γ = 0 vs. H_1_: γ > 0 using Z-test, where γ = 0 (=1) is indicative of non-significant introgression. We applied HyDe to the concatenated supermatrix SuperMatrix_50BUSCO_dna_pasta_ali_trim of 1603 BUSCO genes under default parameters specifying *Ephemera danica* as an outgroup. Under this setup HyDe evaluates all possible taxa quartets. Since HyDe only allows indication of a single outgroup taxon (i.e. *Ephemera danica*), we excluded all quartets that contained the *Isonychia kiangsinensis* outgroup from the HyDe output. Additionally, we calculated Patterson’s *D* statistic [57] for every quartet from the frequency (*p*) of ABBA-BABA site patterns estimated by HyDe as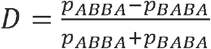 To test significance of *D* statistics we used a *χ*^*2*^ test to assess whether the proportions p_ABBA_ and Pbaba were significantly different. To minimize effect of false positive cases (type I error) in the output, we first applied a Bonferroni correction to the *P* values derived from *Z*- and *χ*^*2*^ tests and then filtered the results based on a significance level of 0.05 and 10^−6^ for *D* and γ, respectively. Also, we noticed that negative values of *D* were always associated with nonsignificant γ values, thus all of the *D* values remained positive after filtering was applied. Additionally, we excluded all quartets that did not match species topology. Further, we ran HyDe on SuperMatrix_50BUSCO_dna_prank_trim with the excluded 3^rd^ codon position to investigate a potential impact of the saturation effect on introgression inference.

*D*_FOIL_ is an alternative site pattern-based approach that detects introgression using symmetric 5-taxon (quintet) trees, i.e. (((P1,P2),(P3,P4)),O). *D*_FOIL_ represents a collection of similar to *D* statistics applied to quintet trees and, if considered simultaneously, they provide a powerful approach to identify introgression including ancestral as well as donor and recipient taxa (i.e. introgression directionality). Moreover, *D*_FOIL_ exhibits exceptionally low false positive rates [129]. Since the number of possible quintet topologies for a phylogeny of 85 taxa is > 32×10^6^, for analysis we extracted them only for every odonate suborder individually with the custom R scripts. Note, that for Anisozygoptera we only considered quintets that can be formed between Anisozygoptera, Anisoptera, and Zygoptera taxonomic groups. As the number of Anisozygoptera quintets is highly disproportional (34619 out of all 72971 tested Odonata quintets), for downstream analyses we randomly selected 4000 Anisozygoptera quintets which approximately matches the number of quintets for an individual species. Analogously to HyDe, we applied *D*_FOIL_ to the concatenated supermatrix SuperMatrix_50BUSCO_dna_pasta_ali_trim of 1603 BUSCO genes under default parameters specifying *Ephemera danica* as an outgroup. Also, since *D*_FOIL_ requires that every quintet has a symmetric topology we considered only those quintets within our phylogeny that met this criterion (Figure 1). Additionally, *D*_FOIL_ requires that the divergence time of P3 and P4 precedes divergence of P1 and P2, i.e. *T*_2_ > *T*_1_, thus we filtered out quintets that violated this assumption using divergence times from our fossil calibrated phylogeny. In order to correct the *P* values resulted from *D*_FOIL_ analysis for multiple testing, we applied the Benjamini-Hochberg procedure at a false discovery rate cutoff (FDR) of 0.05.

As an alternative test for introgression, we performed a simple yet conservative *χ*^2^ goodness-of-fit test on the gene tree count values for each triplet. First, we tested the three count values to see whether their proportions significantly deviate from 1/3, and if the null hypothesis is *not* rejected it will be indicative of extreme levels of ILS. Second, we tested the two count values that originated from discordant gene trees to see whether their proportions significantly deviate from 1/2, and if the null hypothesis is rejected, it will suggest a presence of introgression. In order to correct the *P* values resulted from the *χ*^2^ test for multiple testing, we applied the Benjamini-Hochberg procedure at a false discovery rate cutoff (FDR) of 0.05. Second, we used a branch length test (BLT) to identify cases of introgression [84]. This test examines branch lengths to estimate the age of the most recent coalescence event (measured in substitutions per site). Introgression should result in more recent coalescences than expected under the concordant topology with complete lineage sorting, while ILS yields older coalescence events. Importantly, ILS alone is not expected to result in different coalescence times between the two discordant topologies, and this forms the null hypothesis for the BLT. For a given triplet, for each gene tree we calculated the distance *d* (a proxy for the divergence time between sister taxa) by averaging the external branch lengths leading to the two sister taxa under that gene tree topology. We calculated *d* for each gene tree and denote values of *d* from the first discordant topology *d*_T1_ and those from the second discordant topology *d*_T2_. We then compared the distributions of *d*_T1_ and *d*_T2_ using a Wilcoxon Rank Sum Test. Under ILS alone the expectation is that *d*_T1_ = *d*_T2_, while in the presence of introgression *d*_T1_ < *d*_T2_ (suggesting introgression consistent with discordant topology T_1_) or *d*_T1_ > *d*_T2_ (suggesting introgression with consistent with topology discordant T_2_). The BLT is conceptually similar to the D3 test [130], which transforms the values of *d*_T1_ and *d*_T2_ in a manner similar to the *D* statistic for detecting introgression. As with the DCT, we performed the BLT on all triplets within a clade and used a Benjamini-Hochberg correction with a false discovery rate cutoff (FDR) of 0.05. We note that both the *χ*^2^ test and BLT will be conservative in cases where there is bidirectional introgression, with the extreme case of equal rates of introgression in both directions resulting in a complete loss of power.

QuIBL is based on the analysis of branch length distributions across gene trees to infer putative introgression patterns. Briefly, under coalescent theory internal branches of rooted gene trees for a set of 3 taxa (triplet) can be viewed as a mixture of two distributions with the underlying parameters. Each mixture component generates branch lengths corresponding to either ILS or introgression/speciation. Thus, estimated mixing proportions (π_1_ for ILS and π_2_ for introgression/speciation; π_1_ + π_2_ = 1) of those distribution components show what fraction of the gene trees were generated through ILS or non-ILS processes. For a given triplet, QuIBL computes frequency of gene trees that support three alternative topologies. Then for every alternative topology QuIBL estimates mixing proportions along with other relevant parameters via Expectation-Maximization and computes Bayesian Information Criterion (BIC) scores for ILS-only and introgression models. For concordant topologies elevated values of π_2_ are expected whereas for discordant ones π_2_ can vary depending on the severity of ILS/intensity of introgression. In extreme cases when the gene trees were generated exclusively under ILS, π_2_ will approach to zero and the expected gene tree frequency for each alternative topology of a triplet will be approximately 1/3. To identify significant cases of introgression here we used a stringent cutoff of ΔBIC < -30 [68]. We ran QuIBL on every triplet individually under default parameters with number of steps (`numsteps` parameter) is equal to 50 and specifying one of the Ephemeroptera species for triplet rooting. For computational efficiency we extracted triplets only from the odonate superfamilies in a similar manner as we did for D_FOIL_ (see above). For this analysis we used 1603 ML gene trees estimated from CO orthology clusters.

In order to estimate phylogenetic networks from the 1603 ML gene trees estimated from CO orthology clusters, we used pseudolikelihood (InferNetwork_MPL) and likelihood (CalGTProb) approaches implemented in PhyloNet. For both analyses we only selected gene trees that had at least one of the outgroup species (*Isonychia kiangsinensis* and *Ephemera danica*) and at least three ingroup taxa. We performed network searches for our focal clades, namely Anisozygoptera, Aeshnidae, Gomphidae+Petaluridae and Libellulidae. For pseudolikelihood analysis, we ran PhyloNet on every clade individually allowing a single reticulation event, with the starting tree that corresponds to the species phylogeny (`-s` option), 100 iterations (`-x` option), 0.9 bootstrap threshold for gene trees (`-b` option) and optimization of branch lengths and inheritance probabilities on the inferred networks (`-po` option). To ensure convergence, the network searches were repeated 3 times. For Anisozygoptera, we explicitly indicated *Epiophlebia superstes* as a putative hybrid (`-h` option). For the full likelihood estimation, we fixed the topology of a focal clade (equivalent to the species tree topology) and calculated likelihood scores for possible networks with a single reticulation (generated with a custom script) using CalGTProb. Additionally, to assess significance of networks, we used difference of BIC scores (ΔBIC) derived from network without reticulation (i.e. tree) and a network with a reticulation (Data S6).

### Dimensionality reduction and visualization

To uncover and visualize complex relationships between site pattern frequencies and Patterson’s D statistic and Hyde γ parameter we implemented a dimensionality reduction technique t-distributed stochastic neighbor embedding (tSNE) [65] under default parameters in R. Specifically we estimated tSNE maps from counts of 15 quartet site patterns calculated by HyDe (“AAAA” ,”AAAB” ,”AABA”, “AABB”, “AABC”, “ABAA”, “ABAB”, “ABAC”, “ABBA”, “BAAA” ,”ABBC”, “CABC”, “BACA”, “BCAA”, “ABCD”).

## QUANTIFICATION AND STATISTICAL ANALYSES

All statistical analyses were performed in R (https://www.R-project.org/). Statistical significance was tested using the Wilcoxon Rank Sum test, Fisher Exact test and/or *χ*^2^ test.

## DATA AND SOFTWARE AVAILABILITY

All raw RNA-seq read libraries generated for this study have been uploaded to the Short Read Archive (SRA) (SRR12086708-SRR12086765). All assembled transcriptomes, alignments, inferred gene and species trees, fossil calibrated phylogenies (including paleontological record assessed from PaleoDB used to plot the fossil distribution in Figure 1), PhyloNet results, QS results and introgression results are available on figshare (https://doi.org/10.6084/m9.figshare.12518660). Custom scripts that were used to create various input files as well as the analysis pipeline are available on GitHub https://github.com/antonysuv/odo_introgression.

