## Supplementary Information for "Deep ancestral introgression shapes evolutionary history of dragonflies and damselflies"

**Supplemental Information**

**SI Results**

**Distribution of *D* statistic.**

First, we compared the distributions of *D*-statistic values between (Figure S4A) different taxa computed from ABBA-BABA [1] site patterns (Figure 3A) by applying HyDe to various groups of four species (quartets). Although quartets with older divergence times tend to have only slightly decreased values of *D* [2] (Wilcoxon rank-sum test [WRST], *P* = 3.116🞨10^-8^), this effect can be more pronounced for extremely old divergences. Indeed, quartets that were formed within the most recently diverged clades in our phylogeny, including Calopterygoidea (Late Cretaceous, ~ 67 mya), Coenagrionoidea (Early Cretaceous, ~ 116 mya) and Libelluloidea (Late Cretaceous, ~ 87 mya) have significantly greater *D* values (WRST, *P* = 2.276🞨10^-9^) if compared with quartets that were formed from distantly related species of Zygoptera and Anisoptera (Late Triassic ~237 mya). Quartets involving Zygoptera lineages (i.e. both intra- and inter-Zygoptera comparisons) exhibit significantly higher average *D* statistic values (WRST, all *P* < 0.05, Figure 3A, Data S4) when compared to the distribution of *D* values computed across the entire order. Conversely, Epiprocta, Anisozygoptera, Cordulegastroidea (intra- and inter-superfamilial comparisons) and Libelluloidea (only inter-superfamilial comparisons) showed reduced *D* values (WRST, all *P* < 10^-7^, Figure 3A, Data S4). We compared taxon-specific distributions of significant *D* statistic values to those of the entire order to identify possible extreme skewness in site-pattern frequencies. Overall, we found several mostly minor but significant upward and downward departures of *D* statistic averages from the entire order (Figure S4A). This result may imply that the ILS rate heterogeneity across different odonate linages can differentially affect the value of *D* statistic.

**Distribution of *γ* statistic.**

Overall, comparisons of *γ* distributions (Figure 3A) show several lineages with elevated average *γ* values (WRST, all *P* < 0.05, Figure 3B, Data S4) to the total distribution of *γ*. In contrast to the comparisons of *D* distributions, Epiprocta and inter-superfamilial comparisons of its two groups, Anisozygoptera and Aeshnoidea (Aeshnidae), show significantly higher *γ* (WRST, all *P* < 0.05, Figure 3A, Data S4), whereas all inter-superfamilial comparisons within Zygoptera (except Calopterygoidea), and comparisons within Aeshnoidea (Aeshnidae) and Calopterygoidea show significantly lower *γ* (WRST, all *P* < 0.05, Figure 3A, Data S4).

**Signatures of introgression identified by *D*_FOIL_**

Overall, the distributions of significant *D*_FOIL_ statistics (Figure 4A) except for Anisoptera, introgression scenario 2, Aeshnoidea (Aeshnidae) and Libelluloidea were different from the *D*_FOIL_ distribution within the entire order across all taxonomic levels (WRST, all *P* < 0.05, Figure 4B, Data S5). This observation is indicative of differences in the amount of ancestral introgression as well as its polarization, which is determined by significance of *D*_FOIL_ statistics. Further we note that average *D*_FOIL_ (Figure 4A) was significantly different from 0 (one sample *t*-test [OSTT], all *P* < 0.0223, Figure 4B, Data S5) for all tested cases except Aeshnoidea (Aeshnidae) and Libelluloidea. Together these significant deviations from 0 of *D*_FOIL_ statistics further support the hypothesis of introgression for the tested taxonomic levels [3]. Tests of individual quintets as implemented in *D*_FOIL_ identified significant cases of introgression for all tested introgression scenarios and within all clades except and Aeshnoidea (Gomphidae+Petaluridae) and Lestoidea (Figure 4C).

The analysis of quintet fraction with significant introgression revealed that only introgression scenario 4 and Coenagrionoidea exhibit an excess of significant quintets (FET, all *P* < 0.05, Figure 4C, Data S5), whereas introgression scenario 4 and Calopterygoidea show decrease of significant quintets (FET, all *P* < 0.05, Figure S6C, Data S5) in comparison with the entire order. Despite the notion that *D*_FOIL_ approach exhibits low false positive rate, it requires tree symmetry (Figure 2, see Materials and Methods) [3], thus not all the quintet combinations of taxa can be evaluated for introgression with this method.

**SI Figures**

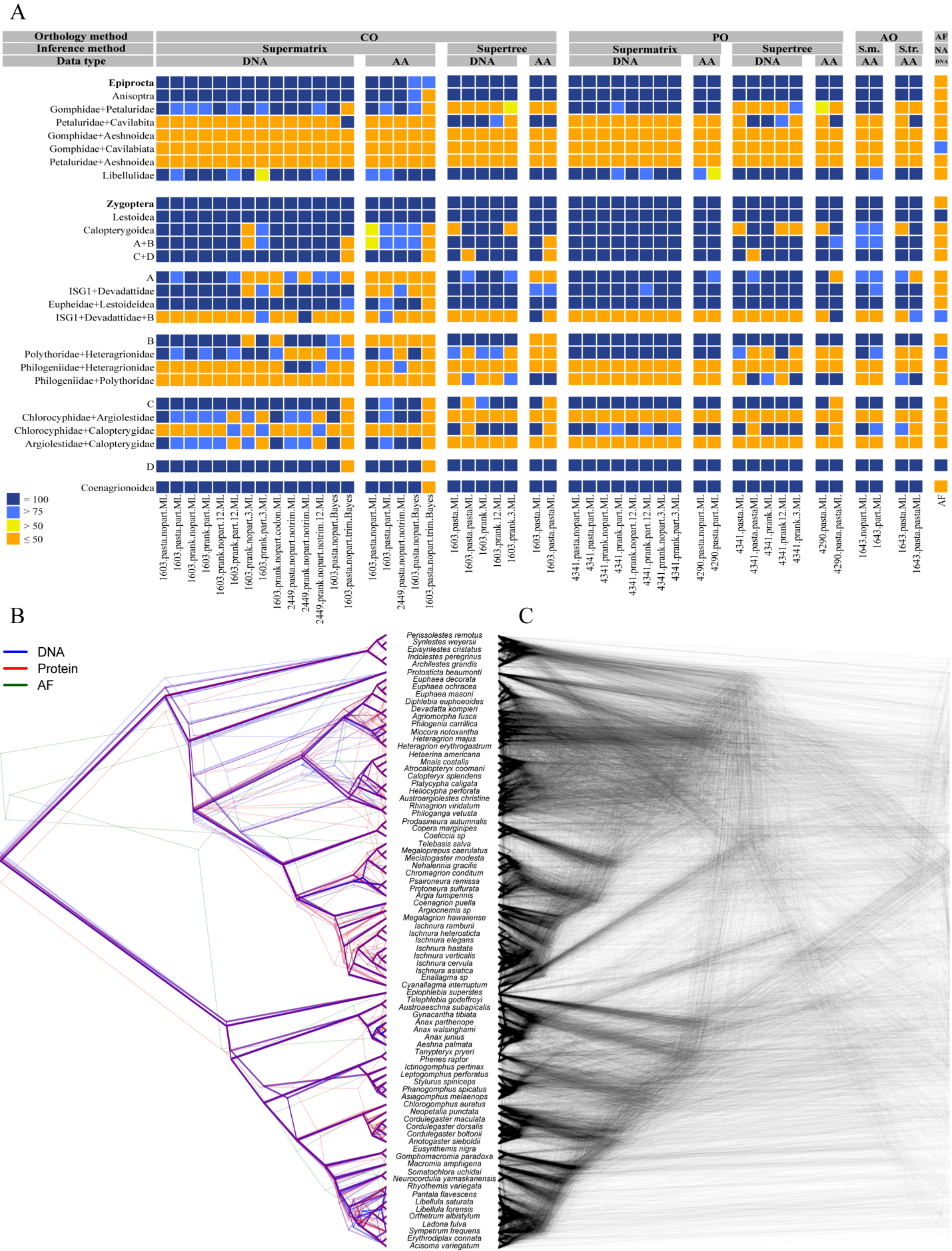
_­_

**Figure S1. Comparison of Orthology Detection and Phylogenetic Tree Reconstruction Pipelines.** (A) Comparison of support for major Odonata group relationship hypotheses. Supports present bootstraps scores for maximum likelihood (ML) and Alignment free (AF) inference, posterior probabilities for Bayesian inference (Bayes) and local branch support from quartet frequencies for Supertree approach using ASTRAL. The text below shows parameters of supermatrix and its analysis types (number_of_loci.alignment_method.partition.inference_method) within Supermatrix framework and parameters for gene alignments and its analysis types (number_of_loci.alignment_method.inference_method) within Supertree framework. The support value ≤ 50 indicates that either insignificant support or these relationships were not observed on a particular phylogeny. CO = single-copy orthologs; AO = all single-copy orthologs; PO = paralogy-parsed orthologs (B) Comparison of phylogenetic tree topologies estimated from different data types. (C) Comparison of BUSCO 1603 gene tree ML topologies.

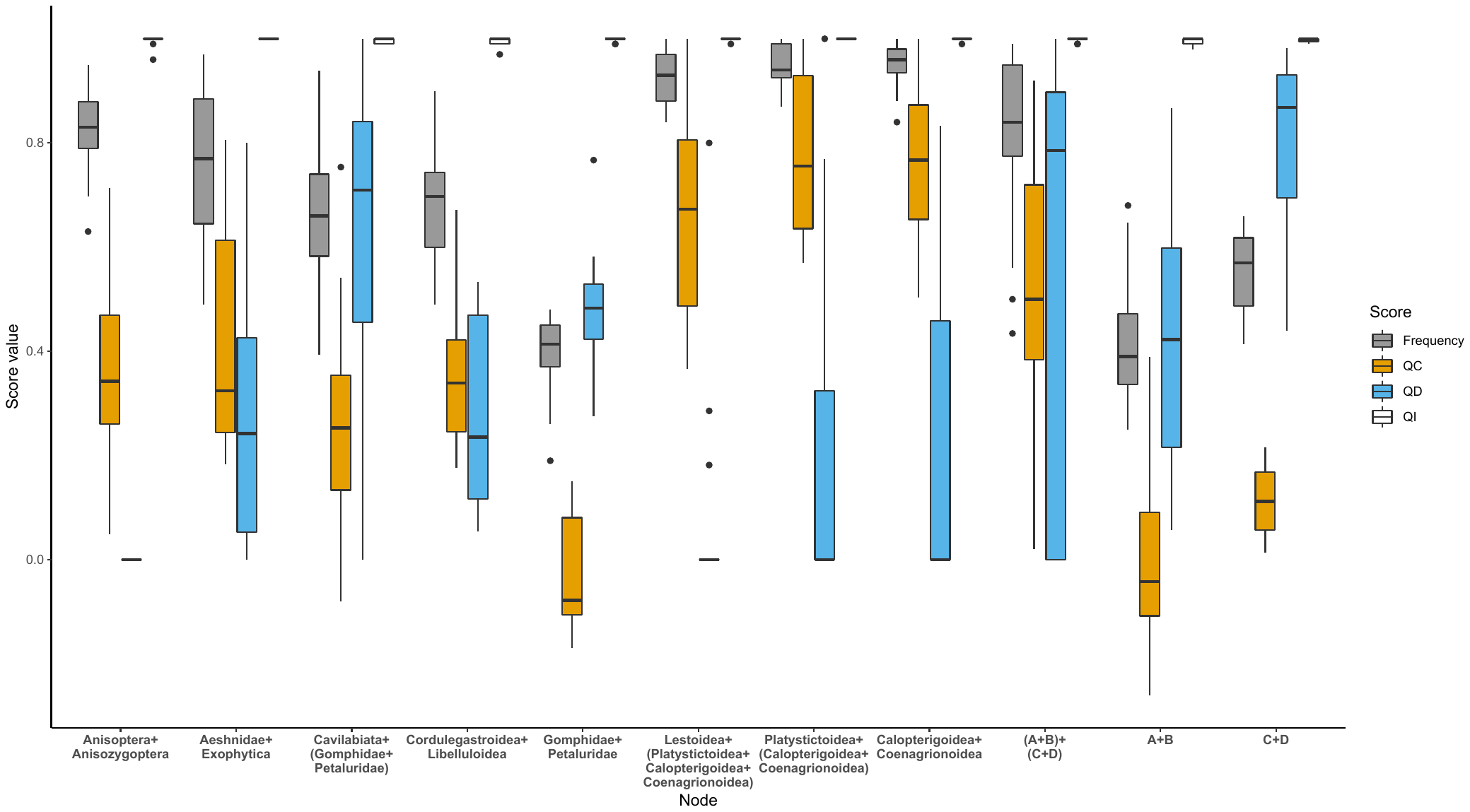

**Figure S2. Quartet Sampling Analysis of Major Odonate Divergence Points.** The boxplots represent distribution of Frequency, Quartet Concordance (QC), Quartet Discordance (QD) and Quartet Informativeness (QI) scores. Frequency indicates proportion of inferred quartets which coincide with the tree topology. QC indicates how often concordant quartets are inferred over the discordant ones. QD shows bias toward any particular discordant quartet. QI indicates whether the quartets are informative or not. All of the scores were derived for 15 supermatrices using tree topology as in Figure 1.

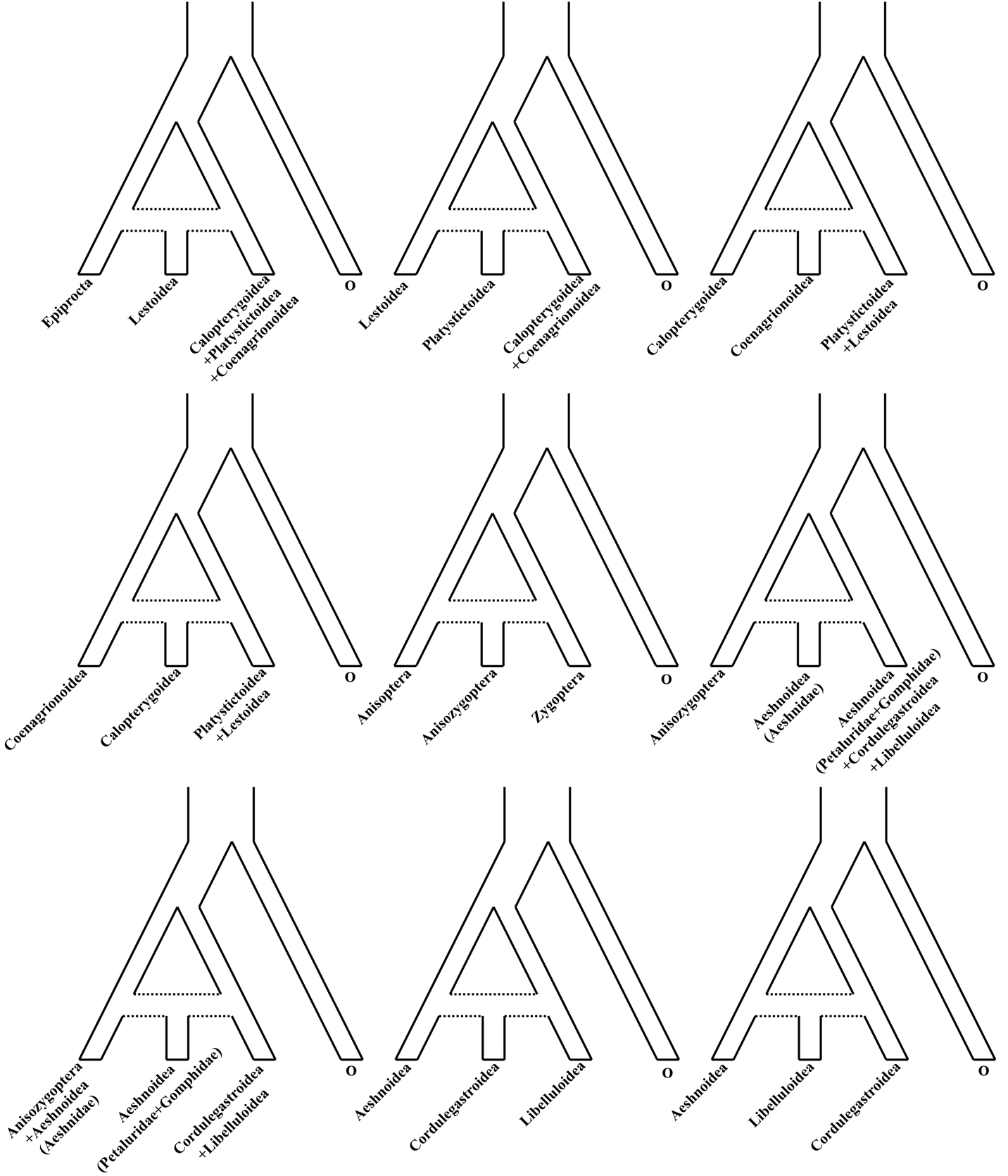

**Figure S3. Hypotheses of Introgression/Hybridization scenarios between Odonata Superfamilies Tested in HyDe using D and** *γ* **Statistics.**

**Figure S4. Distributions of the Patterson’s D Statistic and Relations between HyDe γ and D.**

(A) Estimated distributions of significant (Bonferroni corrected P < 0.05) D statistic from ABBA-BABA site pattern counts for each quartet using HyDe output. Black dots mark medians of violin plots. Asterisks indicate significantly greater (red) or lower (blue) D averages of various tested cases compared to D average of the entire order.

(B) Non-linear relationships between absolute values of significant D statistics and γ. The black line denotes a GAM fit. The color legend shows density of quartets across γ-D plane.

(C)-(D) tSNE projections of the 15 site pattern counts derived for each of the 32620 significant quartets from HyDe output (each dot on a tSNE map represents a quartet). Color schemes reflect significant values of D statistic (E) and significant values of of γ (F).

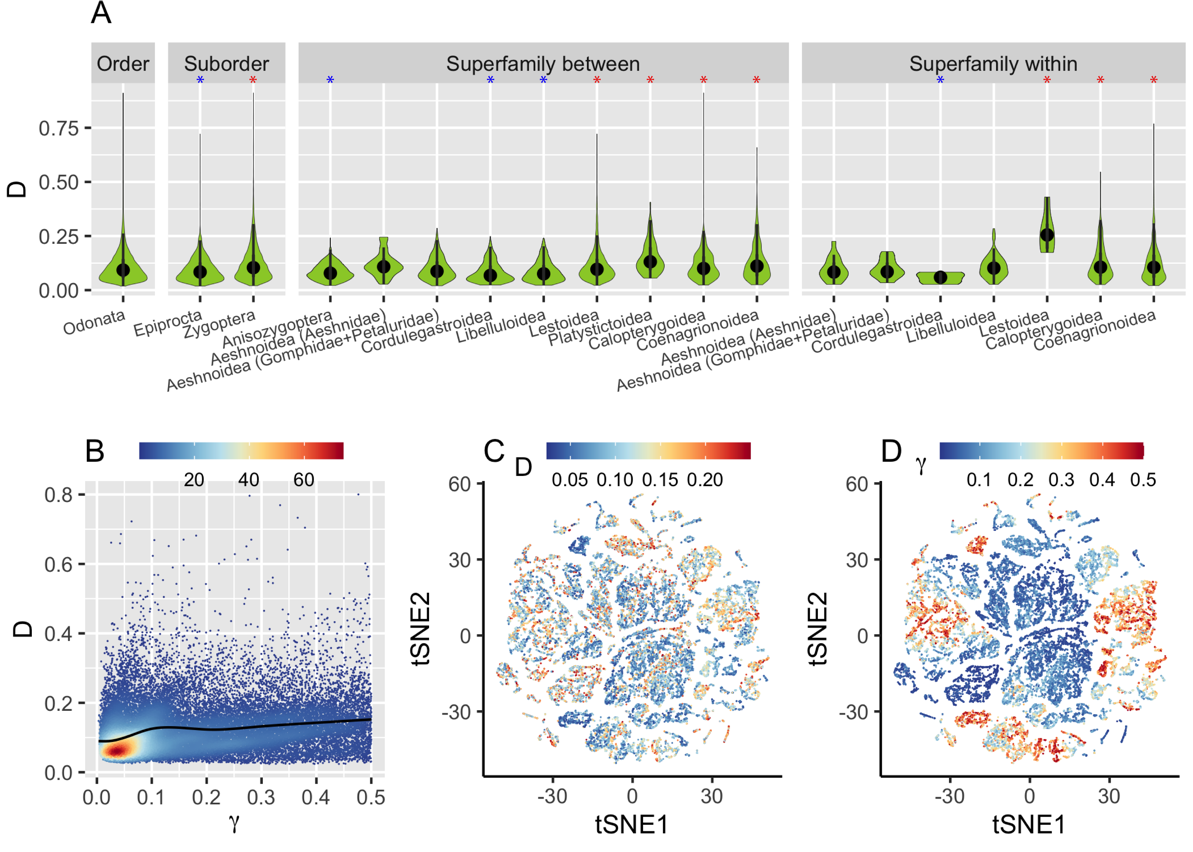

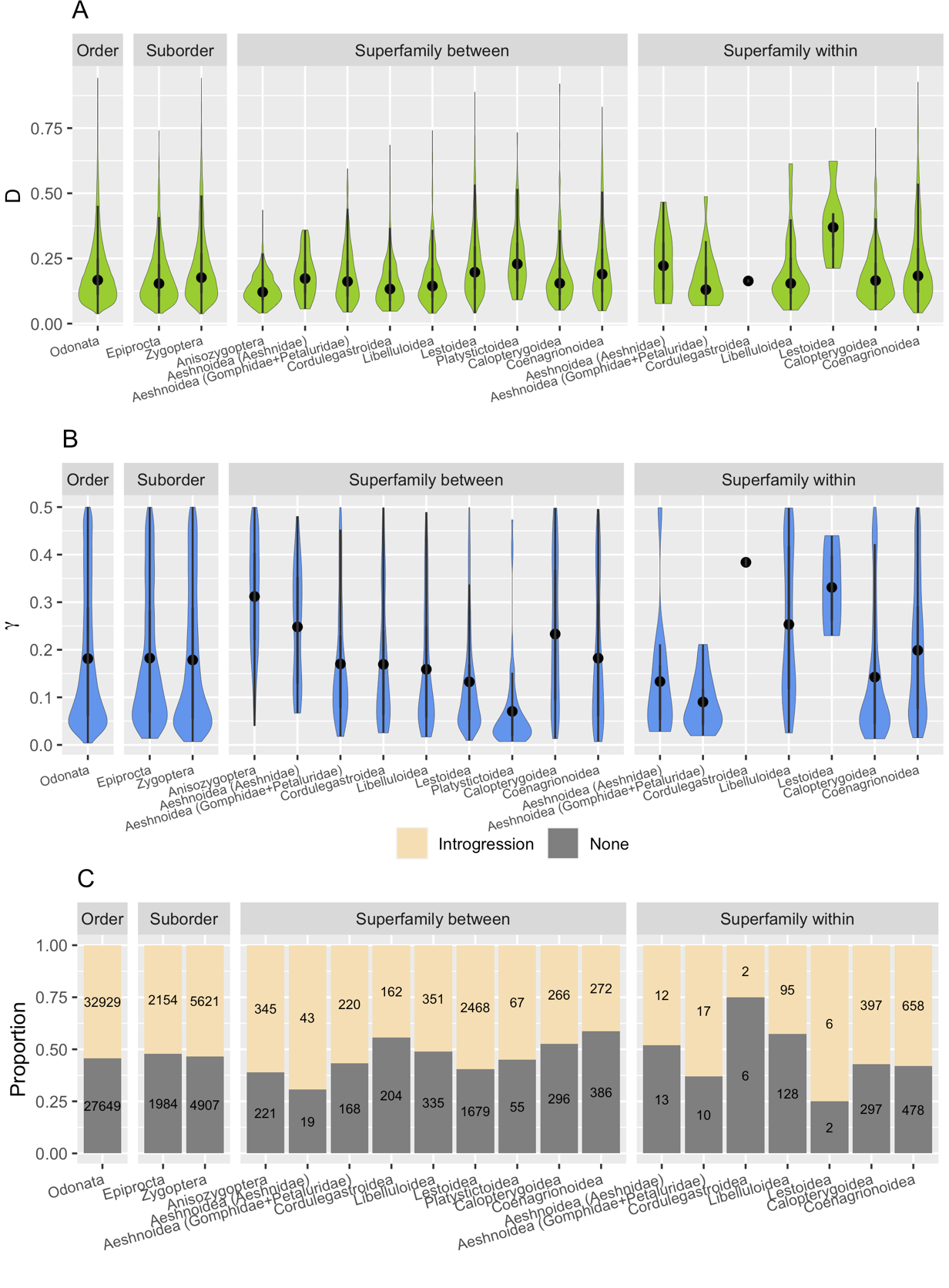

**Figure S5. Distributions of the Patterson’s D Statistic, HyDe γ and Their Relations Across Odonate Taxonomic Levels using the 1^st^ and 2^nd^ codon positions.**

(A) Estimated distributions of significant (Bonferroni corrected P < 0.05) D statistic from ABBA-BABA site pattern counts for each quartet using HyDe output. Black dots mark medians of violin plots.

(B) Distribution of significant (Bonferroni corrected *P* < 10^-6^) γ values for each quartet estimated by HyDe. In general, γ values that are not significantly different from 0 denote no relation of a putative hybrid species to either of the parental species P_1_ (1-γ) or P_2_ (γ) in a quartet.

(C) Proportions of quartets that support or reject introgression based on simultaneous significance of *D* statistics and

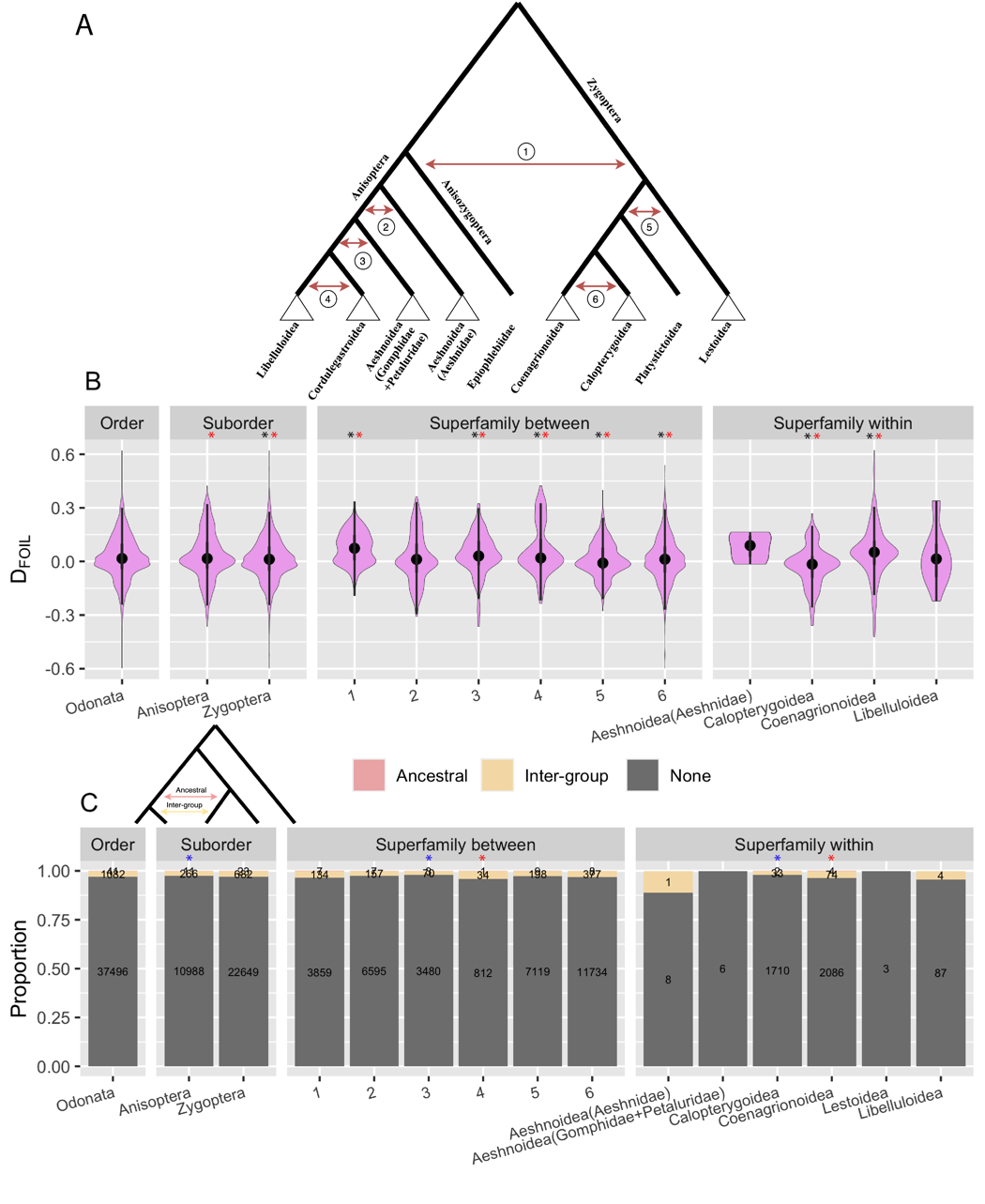

**Figure S6. Distributions of the *D*_FOIL_ Statistics and Their Relations Across Odonate Taxonomic Levels.**

(A) Tested scenarios of deep (numbered red arrows) and intra-superfamilial (white triangles) introgression for Anisoptera, Anisozygoptera and Zygoptera using *D*_FOIL_.

(B) Estimated distributions of *D*_FOIL_ statistics from different site pattern counts for each symmetric quintet. *D*_FOIL_ allows to determine introgression and its polarization between a donor and recipient taxa by comparison of a sign (+/-/0) for all *D*_FOIL_ statistics. Black dots mark medians of violin plots. Lestoidea and Aeshnoidea (Gomphidae+Petaluridae) had no significant cases. Black asterisks indicate significant deviation of *D*_FOIL_ averages of various tested cases compared to *D*_FOIL_ average of the entire order. Red asterisks indicate whether the *D*_FOIL_ averages significantly different from 0.

(C) Counts of quintets that support ancestral, inter-group or no introgression scenarios based on significance of *D*_FOIL_ statistics (FDR corrected *P* < 0.05) and their directionality. The 5-taxon tree shows the difference between ancestral and inter-group introgression. Asterisks indicate significantly greater (red) and smaller (blue) fraction of quartets that support introgression compared to the entire order.

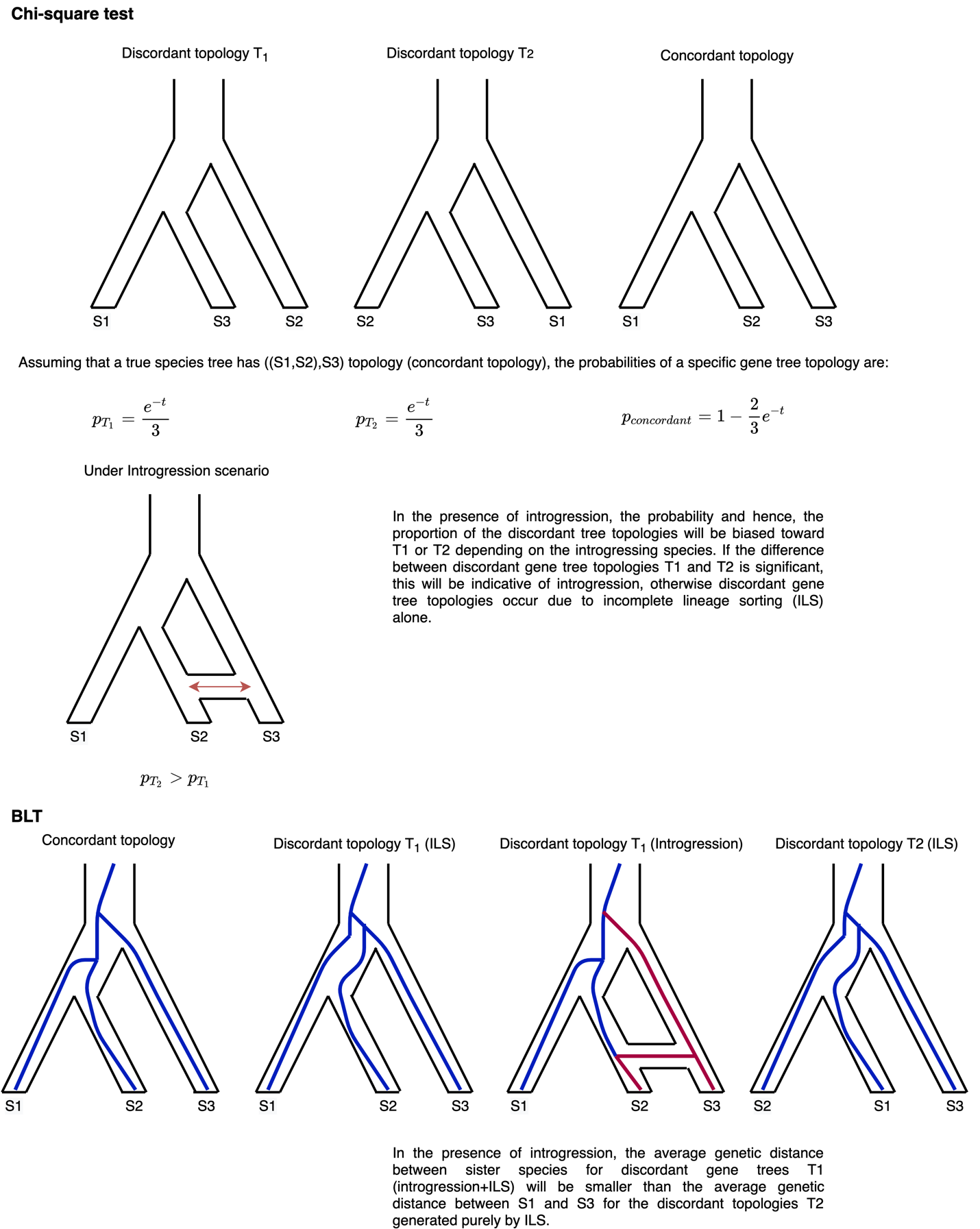

**Figure S7. The Rationale of Chi-square and BLT procedures**

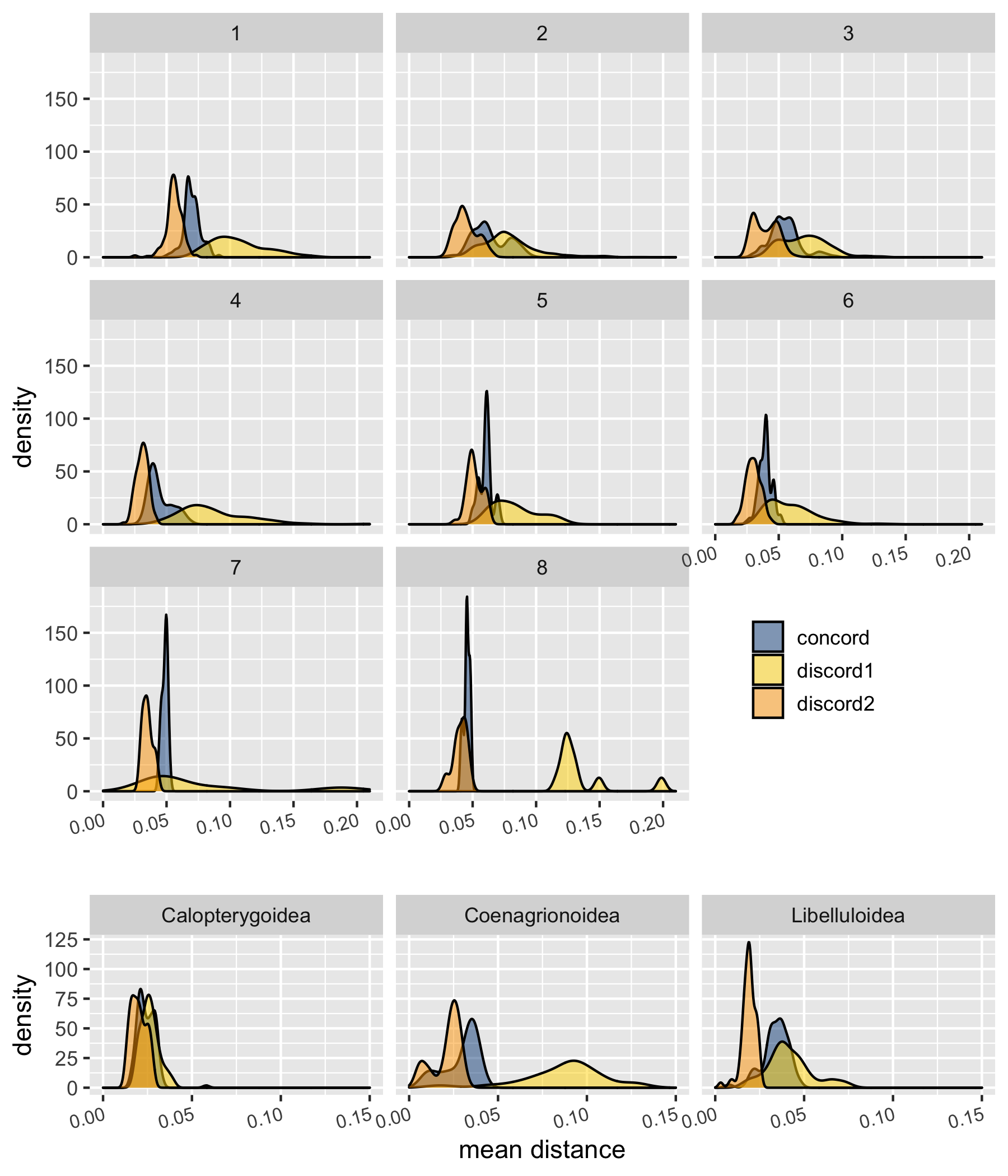

**Figure S8. Normalized genetic mean distance between sister taxa across BUSCO gene trees for concordant and discordant topologies.**

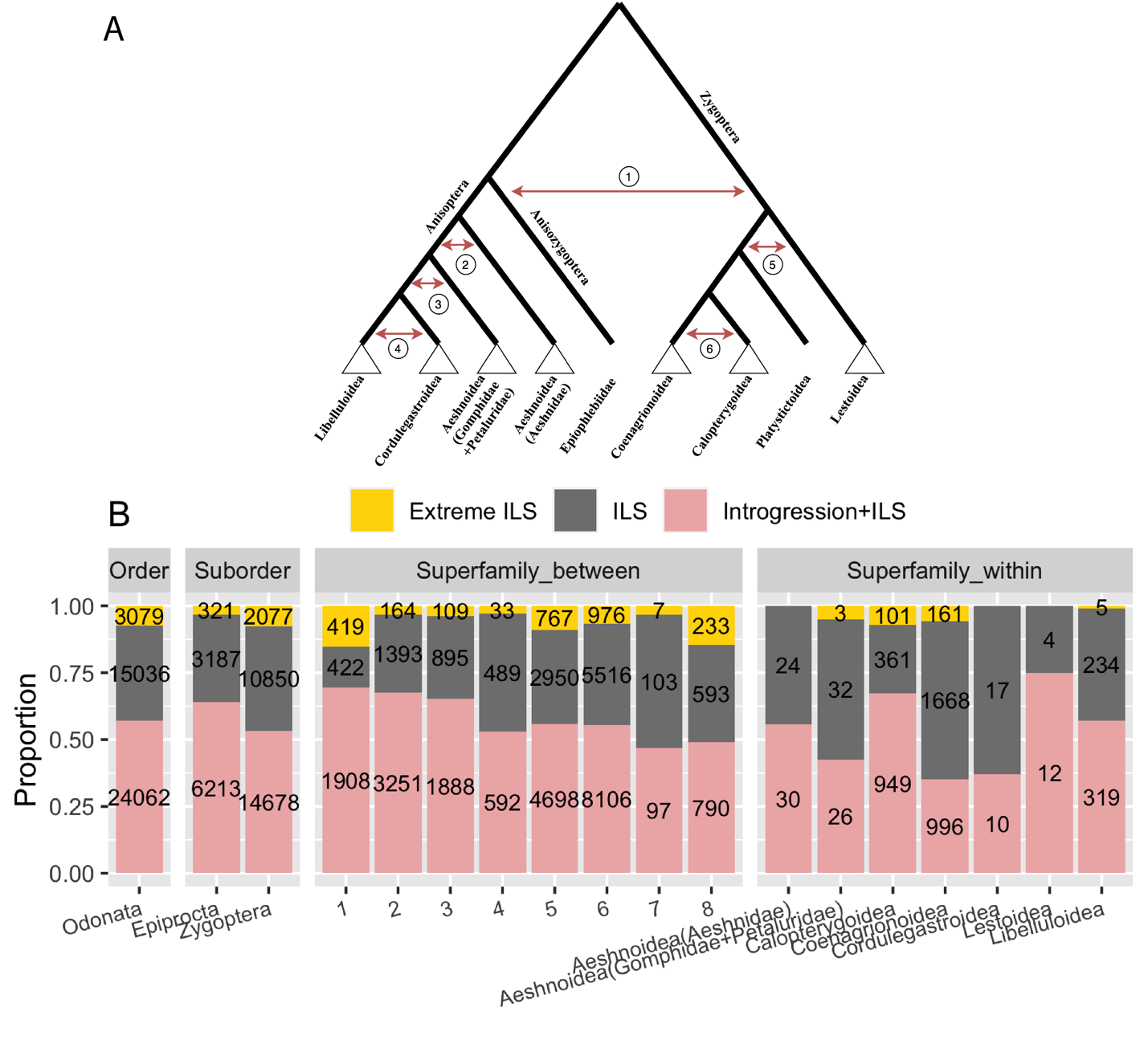

**Figure S9. Overview of QuIBL for Odonate Taxonomic Levels.**

(A) Tested scenarios of deep (numbered red arrows) and intra-superfamilial (white triangles) introgression for Anisoptera, Anisozygoptera and Zygoptera.

(B) Classification of triplets based on BIC criterion. Counts of triplets that are concordant with the species tree (ΔBIC < -30 and maximum number of gene trees agreeing with a triplet), extreme ILS (ΔBIC > -30 and maximum number of gene trees agreeing with a triplet), ILS (ΔBIC > -30) and introgression+ILS (ΔBIC < -30)

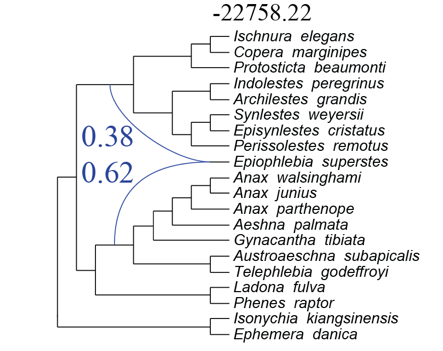

**Figure S10. PhyloNet network inference using pseudo-likelihood approach for Anisozygoptera.**

Phylogenetic network estimated from a set of ML gene trees using pseudo-maximum likelihood approach. *Epiophlebia superstes* was specified as a putative hybrid for Anisozygoptera clade. Blue lines indicate a reticulation event with the value of PhyloNet’s estimated γ. Number above the network indicates the log-likelihood score.

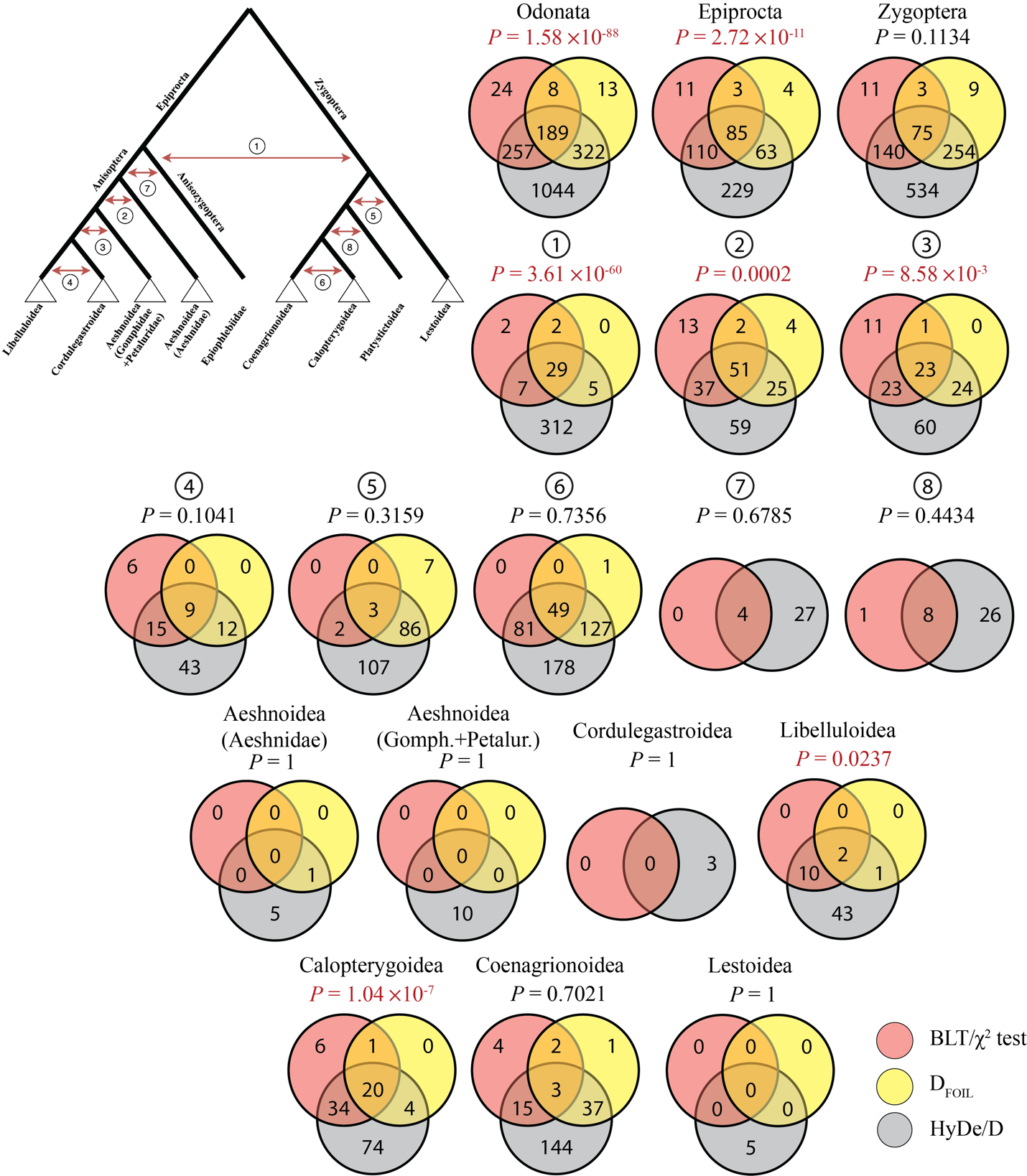

**Figure S11. Overlap between Putatively Introgressed Species Pairs Inferred by Hyde/D, D_FOIL_ and BLT/χ^2^ test**.

The tree shows tested scenarios of deep (numbered red arrows) and intra-superfamilial (white triangles) introgression for Anisoptera, Anisozygoptera and Zygoptera. The numbers within sets represent the number of unique introgressing species pairs identified by a corresponding method. Significance of an overlap between all methods (intersection of all sets) for each scenario was determined by the exact multi-set interactions test. Significant P values are indicated in red. Note that due to the limitations of D_FOIL_, introgression could not be tested for scenarios 7 and 8 as well as within Cordulegastroidea using this method.

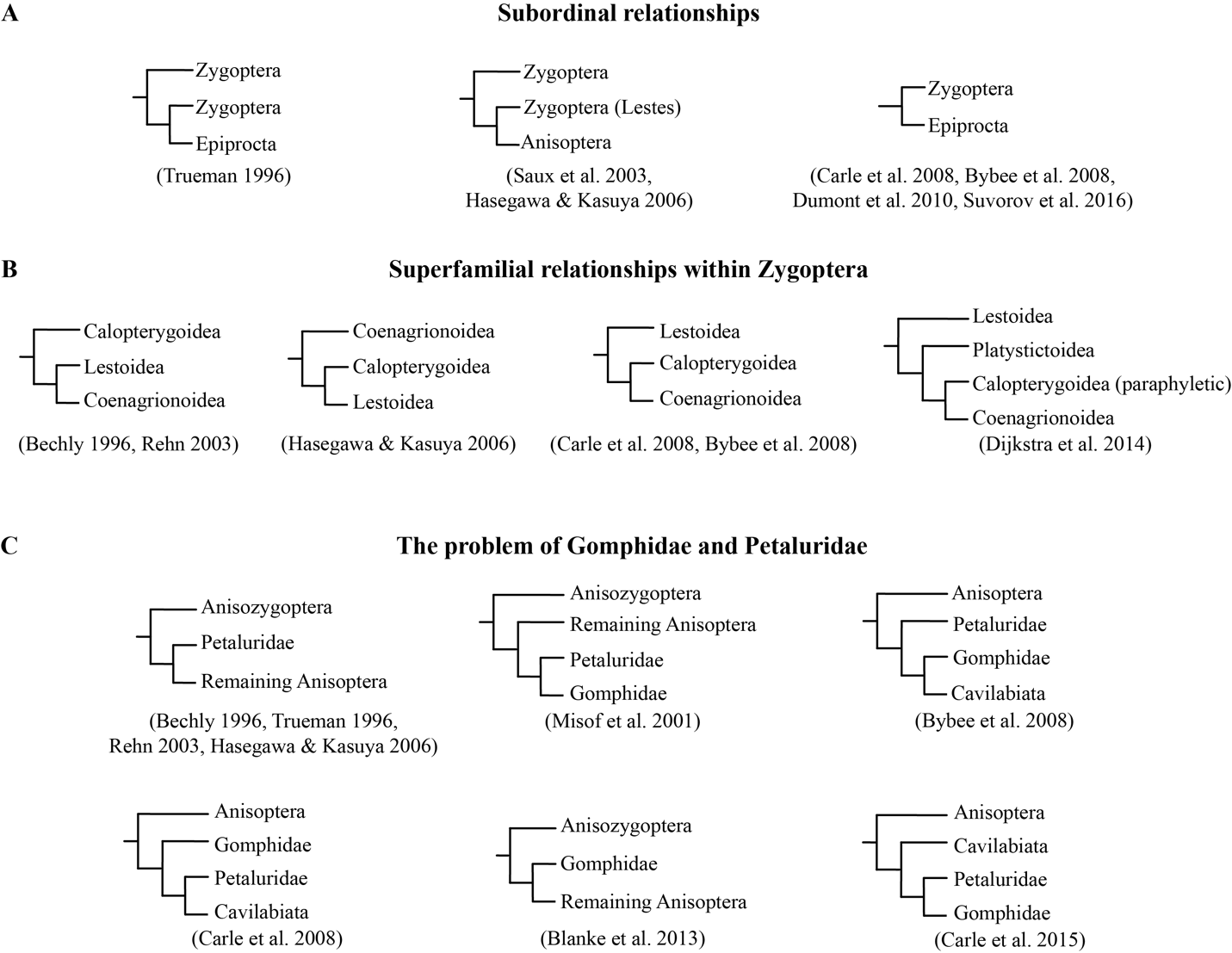

**Figure S12. Phylogenetic Hypotheses of Odonata.**
